# Low-intensity pulsed ultrasound stimulation (LIPUS) modulates microglial activation following intracortical microelectrode implantation

**DOI:** 10.1101/2023.12.05.570162

**Authors:** Fan Li, Jazlyn Gallego, Natasha N Tirko, Jenna Greaser, Derek Bashe, Rudra Patel, Eric Shaker, Grace E Van Valkenburg, Alanoud S Alsubhi, Steven Wellman, Vanshika Singh, Camila Garcia Padill, Kyle W. Gheres, Roger Bagwell, Maureen Mulvihill, Takashi D.Y. Kozai

## Abstract

Microglia are important players in surveillance and repair of the brain. Their activation mediates neuroinflammation caused by intracortical microelectrode implantation, which impedes the application of intracortical brain-computer interfaces (BCIs). While low-intensity pulsed ultrasound stimulation (LIPUS) can attenuate microglial activation, its potential to modulate the microglia-mediated neuroinflammation and enhance the bio-integration of microelectrodes remains insufficiently explored. We found that LIPUS increased microglia migration speed from 0.59±0.04 to 1.35±0.07 µm/hr on day 1 and enhanced microglia expansion area from 44.50±6.86 to 93.15±8.77 µm^2^/min on day 7, indicating improved tissue healing and surveillance. Furthermore, LIPUS reduced microglial activation by 17% on day 6, vessel-associated microglia ratio from 70.67±6.15 to 40.43±3.87% on day 7, and vessel diameter by 20% on day 28. Additionally, microglial coverage of the microelectrode was reduced by 50% in week 1, indicating better tissue-microelectrode integration. These data reveal that LIPUS helps resolve neuroinflammation around chronic intracortical microelectrodes.

## 1. Introduction

Inflammation plays a critical role in our brain’s defense against diseases and damage^1^. Although acute inflammation helps to clear cellular debris, chronic inflammation may exacerbate the tissue damage^2^. Particularly, chronic inflammation impairs neurovascular coupling^3^, disrupting the energy supply to the brain, and elevating oxidative stress^4^. Furthermore, neuroinflammation increases the permeability of the blood-brain barrier (BBB)^5^, allowing infiltration of blood contents such as fibrinogen, which can lead to damage in brain tissue^6^. These changes induced by inflammation are linked to aging process^7^, pain sensation^8^, gliomas^9^, injuries induced by stroke^10^, and traumatic brain injury^11^, as well as neurodegenerative diseases such as Alzheimer’s disease^12^, multiple sclerosis^13^, and Parkinson’s disease^14^.

Neuroinflammation is a complex process orchestrated through interactions of blood cells^15^, endothelial cells^16^, and glial cells^17^. Among glial cells, microglia are first responders and important mediators of neuroinflammation, serving to protect the brain from injury and diseases^18^. Their fine filipodia, or processes, enable them to efficiently survey the surrounding area, detecting pathogens, or disturbances^19^. Microglia express multiple cell surface receptors capable of binding to damage-associated and pathogen-associated molecular patterns^20^, allowing them to quickly respond and migrate to the injury site during the acute phase after an injury (day 0-4)^21^. During this phase, microglia tend to extend both longer and greater number of processes toward the injury site while reducing the length and number of processes away from the injury site^22^. Furthermore, microglia contribute to the resolution of inflammation via clearance of necrotic brain tissue and cellular debris^23,24^. Moreover, they can accumulate around blood vessels to prevent BBB leakage by upregulating tight junction proteins^25^ and promoting vascular remodeling via angiogenesis^26^. Collectively, the evidence strongly suggests that microglial activation can facilitate tissue recovery and safeguard the integrity of the BBB during the acute phase. Though acute microglial activation benefits tissue health, the sustained activation of microglia can exacerbate inflammation. Over days following inflammation, there is a significant proliferation of microglia, leading to microgliosis around the site of injury or disease^27^. During this period, proinflammatory microglia promote the release of nitric oxide (NO)^28^, causing abnormal dilation of the cerebral vessels^29^. Additionally, those microglia can attach to blood vessels, initiate upregulation of proinflammatory profiles, and phagocytose astrocyte endfeet, breaking down the neurovascular unit^25^. Phagocytic microglia also contributes to the loss of neurons and synapses via complement activation^30,31^, disrupting the neural circuit and adversely affecting the propagation of neural signals.

These microglial processes can often be detrimental to intracortical microelectrode interfaces, which can detect neuronal signals and study neural activity^32–35^. Remarkably, these technologies have demonstrated the potential to restore motor and sensory function via BCIs^36–38^ and rely on efficient electrical signal detection from neural tissue. However, implantation of microelectrodes in the brain inevitably causes injury and induces neuroinflammation and microglial activation^39–42^. Previous studies show that microglia direct their processes toward the microelectrode minutes after implantation, followed by microglial migration toward microelectrodes within 6 – 24 hours^41,43–45^. The microglial migration, accompanied by microglial proliferation, leads to the encapsulation of the microelectrode within the first week^41,43–45^, resulting in microgliosis near the injury site^27^. The microglia encapsulation adds an insulating layer on the microelectrode^46^, and increases the impedance of the microelectrode^47^, and the recorded noise floor^48^. Furthermore, persistent microglial activation upregulates production of proinflammatory cytokines^49–51^ which contributes to progressive neurodegeneration, reducing the number of recorded neurons surrounding the microelectrode^52^ and decreasing the number of detectable single-units^53^. Together with other glia cells, such as astrocytes, NG2 glial cells and oligodendrocytes^39,41,54,55^, this neuroinflammatory response increases the noise in electrophysiological recordings and prevents the tissue-microelectrode integration. Although administration of drug such as dexamethasone^42^ and HOE-642^40^ or coating microelectrodes with zwitterionic polymer^56^ and neuroadhesive L1^45^ reduced microglial activation, these interventions require either invasive and recurrent injections or complex manufacturing process^57^. Therefore, modulating microglial changes following microelectrode implantation remains a challenge in neuroscience research and BCI applications.

Ultrasound stimulation is an emerging tool for neuromodulation, achieved through delivery of high-frequency mechanical waves to neural tissue^58–60^. Particularly, low-intensity pulsed ultrasound stimulation (LIPUS) delivers intermittent waves that have been shown to reduce inflammation and promote tissue healing without causing overwhelming thermal effects^61^. Recent studies report that LIPUS can also have a gliomodulatory effect, polarizing microglia toward anti-inflammatory phenotypes^62^, increasing BDNF signaling pathways^63^ and reducing proinflammatory cytokine expression in microglia^64^. These changes in microglia could suppress prolonged inflammation and promote tissue recovery. Therefore, LIPUS has the potential to heal damage caused by microelectrode implantation via attenuation of microglia-mediated neuroinflammation. Compared to administration of drugs and microelectrode coating techniques, LIPUS has the potential to target tissue at the site of injury with high spatial resolution and without the need for complex manufacturing processes to add coating materials^65^. In addition, its non-invasiveness^66^ minimizes the side effects such as infusion related reactions during drug injection procedures, inefficient drug release from electrode housing structures, and delamination between coating materials and microelectrode^57,67^.

In this study, we aimed to investigate the effects of LIPUS on microglial activity and vasculatures. Hence, we used two-photon imaging techniques to observe microglial migration, activation, surveillance, encapsulation over microelectrode, density and association with blood vessels. We also monitored changes in diameter of cerebral blood vessels surrounding implanted microelectrodes. We hypothesized that LIPUS treatment could reduce microglial activation, percent microglial coverage on the probe, and the ratio of microglia associated with blood vessels. To test this hypothesis, we employed two-photon microscopy to compare the activity of microglia in a group treated with LIPUS to an untreated control group for a period of 28 days following microelectrode implantation. Our results demonstrated that LIPUS treatment effectively enhanced microglial migration and surveillance. Furthermore, it advanced the microglial transition from activated state to ramified state. In addition, LIPUS reduced the extent of microglial coverage on the probe, the ratio of microglia associated with blood vessels, and the dilation of cerebral blood vessels. These findings suggest that LIPUS has the potential to serve as a noninvasive therapeutic intervention for modulating neuroinflammation, thereby facilitating recovery and improving device-tissue integration following microelectrode implantation.

## 2. Method

### 2.1 LIPUS treatment

#### 2.1.1 Equipment to generate LIPUS

LIPUS was generated by a function generator (135V, Keysight, Santa Rosa, CA, USA), a power amplifier (RF Power Amplifier 2200L, E&I, Rochester, NY, USA) and a 1.13 MHz, single-element focused transducer (#1219, 851 Material, Wrap Electrode, American Piezo, Mackeyville, Pennsylvania) with 22-ms burst lengths at a 4.4% duty cycle and repetition frequency of 2 Hz. The spatial peak temporal average intensity (ISPTA) measured using a hydrophone over the plane transducer in degassed water was 500 mW/cm^2^. To direct the acoustic beam to the targeted region (visual cortex) of the brain, the transducer was positioned using a stereotaxic apparatus.

#### 2.1.2 Acoustic coupling medium for LIPUS stimulation

In order to prevent air from interrupting the wave path, a polyvinyl alcohol (PVA) cryogel was applied as an acoustic coupling medium for LIPUS^67^. To prepare the acoustic coupling medium, the PVA powder (PVA: U228-08, Avantor, Center Valley, PA) was subjected to a passive melting process in degassed water at a concentration of 1g of PVA per 10 ml of water. Melting was achieved by placing the PVA powder in a capped container to minimize evaporation. Subsequently, the container was immersed in a water bath to control the temperature below 100℃. The resulting solution, consisting of PVA and water, was then carefully dispensed into cone shaped molds with geometry optimized for focused delivery of LIPUS. The solution solidifies and strengthens through two consecutive freeze-thaw cycles. Each cycle consisted of freezing the molds at -20℃ for 10 hours and then thawing at room temperature for 14 hours. Finally, the PVA cones formed from the molds were stored in degassed water at room temperature. This storage environment ensured the preservation of the required properties for the acoustic coupling medium.

#### 2.1.3 Validation experiments for evaluation of LIPUS power and potential thermal effects

To validate LIPUS power for *in vivo* studies, a submersible hydrophone preamplifier was placed in Kwik-Sil beneath the coverslip and PVA cone to evaluate the delivered power of LIPUS treatment using the paradigm designed in previous sections. The distance between the transducer and hydrophone was set to 14 mm to allow enough space for the PVA cone (Fig. 1a). The attenuation of ultrasound power was assessed in both the X-Y plane (Top right figure in Fig. 1a) and Y-Z plane (Bottom right figure in Fig. 1a). The degree of attenuation was quantified by comparing the power at the various locations to the power at the center of X-Y plane, specifically at the putative brain surface beneath the coverslip. Actual power was estimated to be approximately 300 mW/cm^2^ due to power losses when propagating through the acoustic medium, coverslip, and Kwik-Sil. To estimate the potential thermal effect, a thermocouple (K-style, connected to an Omega TC-08 sensor) was implanted 1.5 mm deep into cortex with LIPUS stimulation over one hemisphere. A total of 15 minutes of LIPUS exposure was performed by maintaining the transducer’s supply voltage at 135 V, divided into three intervals of 5 minutes each, with a 5 minutes’ non-exposure time between each sonication (red and blue periods in Fig. 1b). This sonification paradigm ensured that the temperature change was below 1°C. These stimulation parameters were carefully selected and verified to ensure that the LIPUS could penetrate the coverslip window, effectively delivering the appropriate level of stimulation to the tissue surrounding the microelectrode implant.

**Figure 1.**
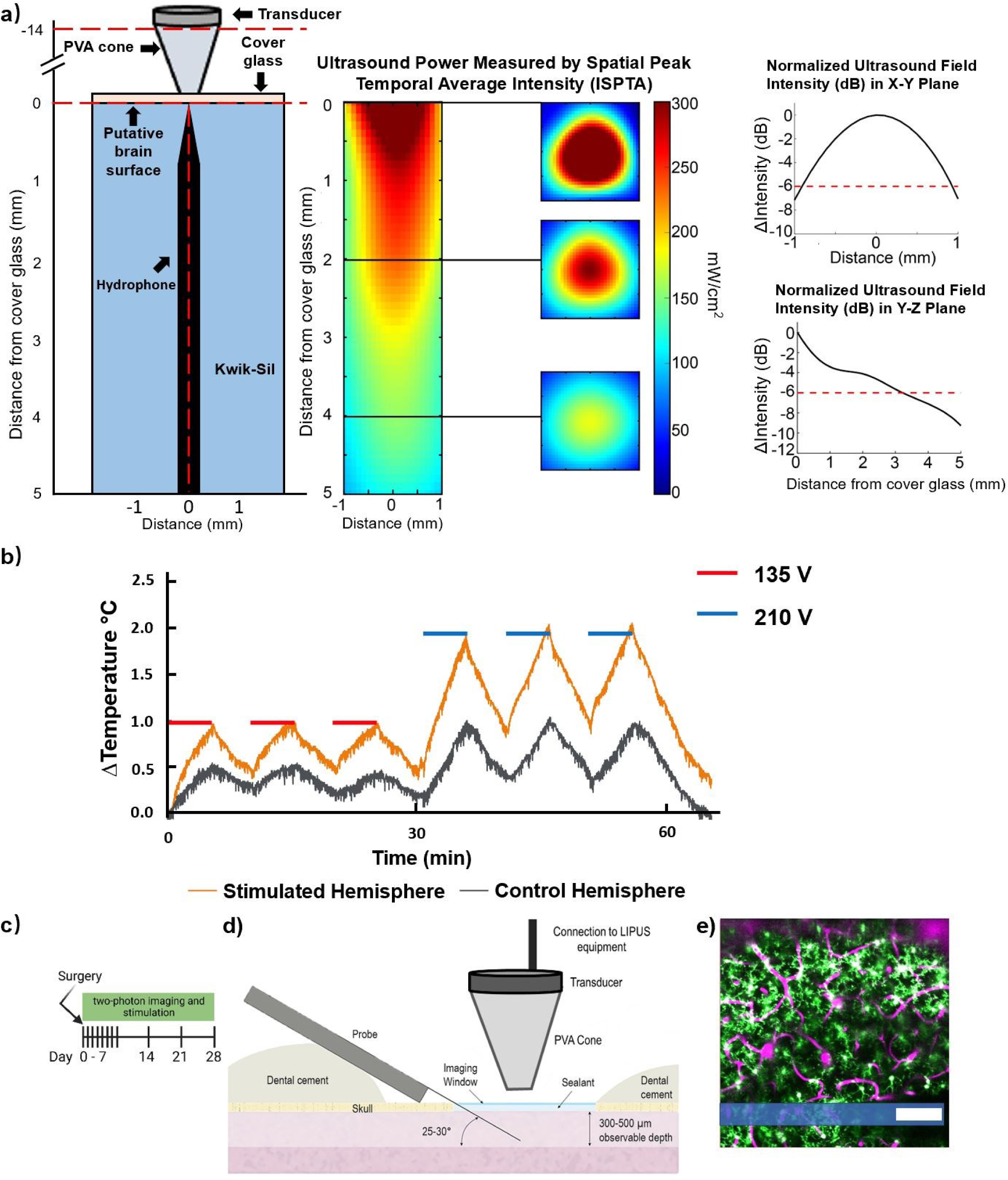
Experimental setups for LIPUS treatment and two-photon imaging of microglia and vasculature following microelectrode implantation. **a)** A submersible hydrophone preamplifier was placed in Kwik-Sil (blue area) below the coverslip and PVA cone to evaluate the LIPUS stimulation power. The measurement of the spatial-peak temporal-average intensity (ISPTA) was conducted by setting the transducer’s voltage supply to 135 V. The power was around 300 mW/cm^2^ at the center of the plane, 1 mm below the cover glass. The imaging planes covered a depth range from 0 to 0.3 mm below the cover glass. The attenuation of ultrasound power was assessed in both the X-Y plane (Top right figure) and the Y-Z plane (Bottom right figure). The degree of attenuation was quantified by comparing the power at the various locations to the power at the center of X-Y plane, specifically at the putative brain surface beneath the coverslip. **b)** Quantification of thermal effect from LIPUS stimulation. By maintaining a power supply voltage of 135 V for the transducer, a total of 15 minutes of LIPUS exposure, divided into three intervals of 5 minutes each, with 5 minutes of non-exposure between each sonication (red and blue periods) can ensure that the temperature change was below 1°C tested at 1.5mm below the cortical surface. **c)** Illustration presents a timeline depicting the sequence of surgery, stimulation, and two-photon imaging. **d)** Schematic representation of microelectrode implantation, sealing of craniotomy window and LIPUS stimulation. **e)** A representative two-photon image of Cx3CR1-GFP transgenic mouse following an I.P. injection of sulforhodamine 101 (SR101) showing microglia cells in green and cerebral blood vessels in magenta. The shaded blue region indicates the location of implanted microelectrode. Scale bar = 100 µm.

### 2.2. Animal preparation

Transgenic CX3CR1-GFP mice (6-8 weeks, N=7 LIPUS group, N=7 control group), which express green fluorescent protein (GFP) under the CX3CR1 promoter in microglia, were used for this study (Jackson Laboratory, Bar Harbor, ME). Each mouse was implanted with a single shank non-functional microelectrode (NeuroNexus, Sample CM16LP). Post-surgery, mice were housed individually to minimize risk of damaging their implant or headcap.

### 2.3. Surgery and probe insertion

Surgical methods employed in this study were consistent with previously described experiments^22,40,41,68^. Anesthesia was induced in mice using a drug cocktail consisting of 75 mg/kg ketamine and 7 mg/kg xylazine, administered intraperitoneally. To secure the anesthetized mice throughout the surgery, a stereotaxic frame (Narishige) was used. Throughout the procedure, the depth of anesthesia was monitored by observing breathing and toe-pinch responses with additional ketamine doses (40 mg/kg) administered hourly or as needed. The animal scalps were shaved and thoroughly washed with betadine and ethanol before being removed along with the connective membranes to expose the skull. Small amounts of Vetbond (3M) were applied to dry the skull surface and enhance the adhesion between the skull and the dental cement headcap. Both the LIPUS-treated mice and control mice underwent chronic surgical procedures. Four bone screws (two over both motor cortices and two over the edges of both visual cortices) were placed and secured to the skull with light-curable dental cement to provide support for a dental cement head cap before craniotomy. A 4x4 mm craniotomy was performed, centered over the ipsilateral visual cortex where the microelectrode was implanted. Throughout the drilling process, saline solution was used to clear bone fragments and maintain a cool surgical site to prevent thermal damage to the brain. After the craniotomy, non-functional single shank silicon probes (NeuroNexus, Sample CM16LP) were inserted at a speed of 200 µm/s through the intact dura mater at a 30-degree angle parallel to the midline, reaching a depth of approximately 300 μm below the pial surface (Fig. 1b). The probe positioning on the brain surface was carefully chosen to avoid major blood vessels. The cranial window was sealed using an in situ curing silicon elastomer (Kwik-Sil, World Precision Instruments) and a glass coverslip, which provided a chronic imaging window for two-photon imaging. The imaging window was secured to the skull using light-curable dental cement, and a 2-mm-high well was built up around the cranial window, to accommodate the water-immersive objective lens used during two-photon imaging. The University of Pittsburgh, Division of Laboratory Animal Resources, and Institutional Animal Care and Use Committee approved all procedures and experimental protocols in strict adherence to the standards for humane animal care as established by the Animal Welfare Act and the National Institutes of Health Guide for the Care and Use of Laboratory Animal.

### 2.4 Two-photon imaging

A two-photon scanning laser microscope (Ultima IV; Bruker) was employed to capture images of CX3CR1-GFP transgenic mice expressing GFP in microglia (Fig. 1c). The microscope setup included a scan head, an OPO laser (Insight DS+; Spectra-Physics), non-descanned photomultiplier tubes (Hamamatsu), and a 16X, 0.8 numerical aperture water immersion objective lens (Nikon). To enhance vascular contrast, mice were injected intraperitoneally with sulforhodamine 101 (SR101). The microscope laser was set to 920 nm to excite both GFP and SR101, and the resulting fluorescence was recorded in the green and red channels, respectively. Z-stack images and ZT-stack images were acquired at various time points post-implantation, including day 0-7, 14, 21, and 28. Throughout all imaging sessions, mice were securely positioned in the stereotaxic setup. The image stacks covered a horizontal area of 412.8 by 412.8 μm (1024 by 1024 pixels), with depths of approximately 300 μm. Images were captured above the top shank and/or below the bottom shank, depending on visibility, which could vary due to the presence of blood vessels or surface bleeding on the pial surface. Only microglia located outside the outer shanks and within the same plane as the probes were included in the subsequent analysis.

### 2.5 Data analysis

The data analysis of images was conducted using ImageJ software (National Institutes of Health)^69^. Specifically, Z-stacks were processed and analyzed to quantify microglial migration, encapsulation, and the blood vessel diameter, of the probes. To further analyze temporal characteristics of microglia activity, ZT-stacks were processed and analyzed to quantify microglial activation, surveillance, and density.

#### 2.5.1 Velocity of migrating microglia

In order to track soma migration velocities, we carefully selected images from Z-stacks captured at consecutive time points to maintain consistency in the region of interest. These images were then used to identify the same microglia throughout the time series as follows. First, the ’TurboReg’ plugin for ImageJ was used to correct the x- and y-axis offsets between a pair of images from two consecutive time points (e.g. day 1 and day 2)^70^. These adjusted images were then merged into a single image stack, with the magenta channel representing the earlier time point and the green channel representing the later time point (Fig. 2a). The time interval between these two time points was recorded as D𝑡. For each image stack, the shortest distance (D𝑥) was measured between the microglia in the magenta channel and their corresponding counterparts in the green channel, with each microglia considered individually. Migration velocity was subsequently calculated using the following formula:

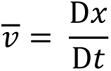

**Figure 2.**
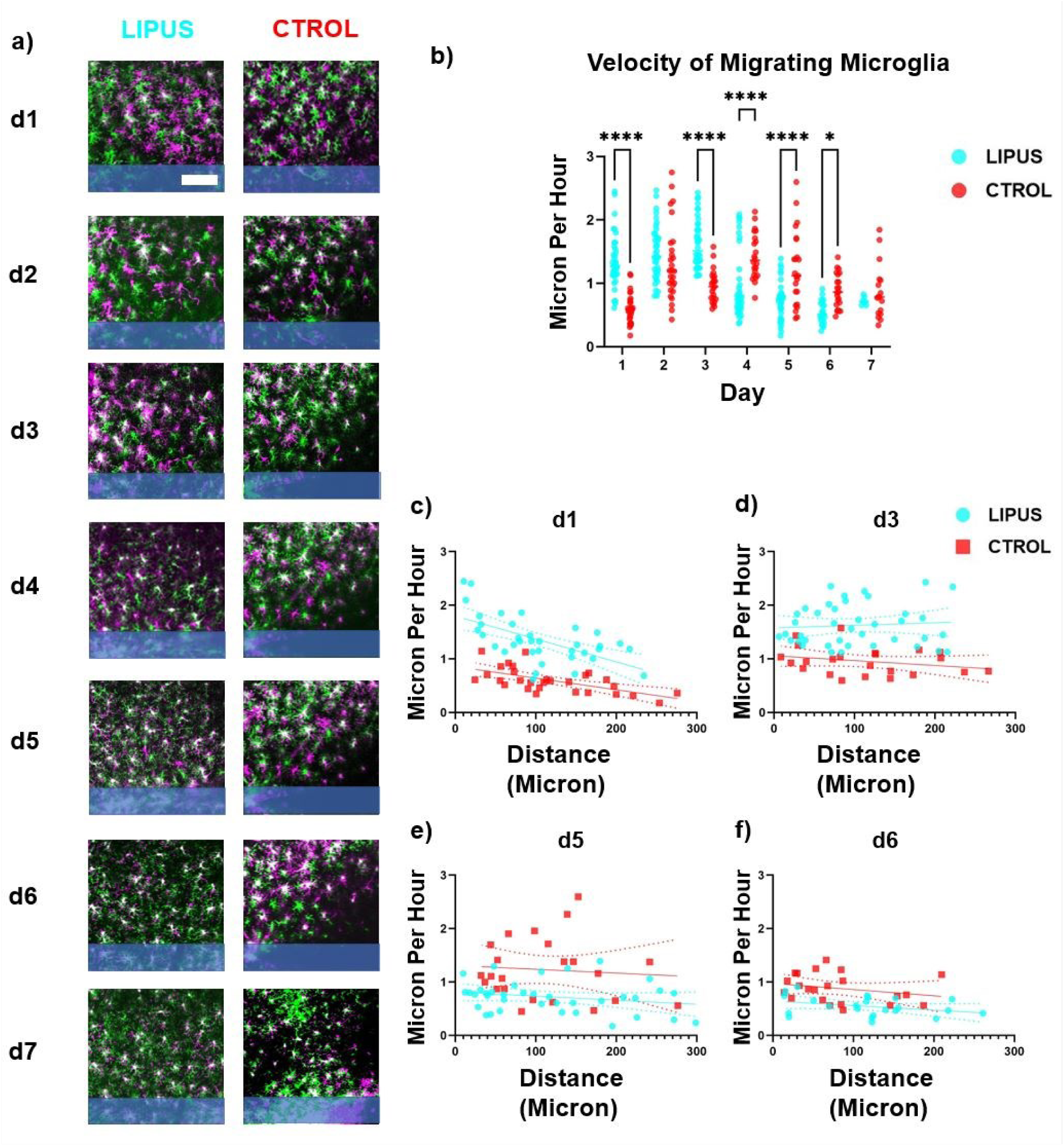
LIPUS increased the velocity of migrating microglia on day 1 and day 3. **a)** Microglia migration was characterized by aligning images from an earlier time point (magenta) and later time point (green). White indicates no cell movements. After microelectrode implantation, microglia migrated toward the microelectrode (shaded blue) as indicated by green cells being closer to the probe compared to magenta cells. **b)** The velocity of migrating microglia was quantified. LIPUS treatment significantly increased the velocity of migrating microglia on day 1 and day 3 but significantly decreased on day 4, 5, 6 (Šídák’s multiple comparisons test, * *p*<0.05, **** *p*<0.0001). **c-f)** spatial characterization of migrating microglia velocity on day 1, 3, 5, and 6. Each microglia was fitted to a linear regression model to examine the relationship between microglia migration velocity and the distance from microelectrode. Significant difference was detected in the intercepts from linear regression model (*p*<0.001) on day 1. Scale bar = 100 µm.

where 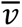 represents the average velocity of the microglia between two time points captured in the images. At each time point, we also recorded the minimum distance from the microglia soma to the manually labeled edge of the probe for the microglia’s specific location.

#### 2.5.2 Microglial activation and morphology

Microglial activation was assessed using a ramification index, which measures the morphological changes in microglia upon activation. When activated, microglia transition from a ramified state to a state with fewer overall processes, but longer processes toward the implantation site^71^. In the image stack, microglia were identified and visually classified as either ramified (1) or transitional (0) microglia in the planes with the probe shank. The distance of these microglia from the surface of the probe was also measured using the ’Measure’ feature in ImageJ, and they were grouped into bins of 50 µm increments up to a distance of 300 µm. For each time point, the data was fitted with a logistic regression to determine the probability distribution of microglia being in the ramified or transitional state as a function of the distance from the probe shank^22,45,68^. The suitability of fitting microglial ramification with a logistic regression model is determined by the receiver operating characteristic (ROC) curve^72^.

In addition to the ramification index, two other indices were calculated to assess microglial activation: the transitional index (T-index) and the directionality index (D-index). The T-index is based on the length of the longest process extending toward (*n*) and away (*f*) from the probe, while the D-index is derived from the number of processes extending toward (*n*) and away (*f*) from the probe. To determine the direction of process extension, a line parallel to the edge of the probe and passing through the midpoint of each microglial soma was used to distinguish the hemisphere toward and away from the implant. Both indices were calculated at each time point using the following formula^22,45,68^:

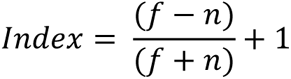

Note that this formula restricts the index values to be zero or positive. In cases where there are more and longer microglial projections directed toward the probe, the value of n (representing the length of processes toward the probe) will be greater than f (representing the length of processes away from the probe), resulting in indices approaching zero. On the other hand, when the processes are evenly distributed in terms of both position and length, n and f will be approximately equal, causing the indices to approach one. Therefore, indices of zero indicate activated microglia, while indices of one indicate a ramified state. However, unlike the binary ramification index, the T-index and D-index also provide additional information about the extent of activation from a morphological perspective.

#### 2.5.3 Microglial surveillance

Microglial surveillance were quantified based on previous literature^73^. To create a series of 2D image projections from 0 to 10 minutes after implantation, grouped z-projections were generated from ZT-stacks with the parameters set to ’Average Intensity’ and a group size of 11, transforming the ZT-stacks into T-stacks. To correct for motion, the ’StackReg’ plugin for ImageJ was applied to align the T-stacks^70^. The images were then thresholded using the ’Li’s Minimum Cross Entropy’ method^74–76^ for better visualization of microglial processes. Microglia surveillance area was measured every minute for 10 minutes. Areas where the GFP signal appeared or disappeared compared to the previous minute were defined as ’expansion’ and ’retraction’, respectively. Two datasets were generated from the thresholded datasets: one dataset included 9 consecutive images from the1^st^ to 9^th^ minute, and the other dataset included images from the 2^nd^ to the 10^th^ minute. The ’subtraction’ operation of the image calculator in ImageJ was used to subtract one dataset from the other and vice versa, resulting in two datasets representing surveillance area expansion and retraction. The average surveillance area expansion and retraction per minute were calculated by counting the average numbers of pixels over 9 images in each stack. The total surveillance area was defined as the area where the GFP signal (from a single microglia) appeared or disappeared at least once within the 10-minute period. The total expansion area and total retraction area were created using the Z-projection feature in ImageJ with the projection type set to ’Max Intensity’. By using the ’OR’ operation of the image calculator in ImageJ, pixels in either the total expansion area or the total retraction area were included in the total surveillance area (blue region in Fig. 4 bottom row). The stable area was defined as the area where the GFP signal (from a single microglia) remained above the threshold throughout the entire 10-minute period. Stable areas (white region in Fig. 4 bottom row) were generated using the Z-projection feature in ImageJ with the projection type set to ’Min Intensity’.

#### 2.5.4 Microglial density

An average of the T-stacks, which were generated in the previous steps, was generated using the ’ZProjection’ feature in ImageJ with the projection type set to ’Average Intensity’. The number of microglia was manually counted within the visible regions, excluding the microelectrode region. The density of microglia was then calculated by dividing this number by the area of the visible region, with the microelectrode area excluded.

#### 2.5.5 Vascular morphology

Vasculature changes were quantified by measuring blood vessel diameter through a Z-stack projection of the entire blood vessel volume. VasoMetrics ImageJ plugin was used for blood vessel diameter, while a skeletonize ImageJ macro was used to quantify tortuosity and branching number and length^77,78^. The vessel coverage was calculated by the vessel area divided by the total area excluding the microelectrode area.

#### 2.5.6 Probe coverage

Microglial encapsulation of the probes was quantified as the percent surface coverage of microglia (GFP) signal^45,79^. The z-stacks were resliced and rotated by 30° using the interactive stack rotation plugin in ImageJ to ensure the entire probe surface was visible in a single frame. The probe surface and the tissue volume up to 20 µm above it were separated into a substack, which was then subjected to a sum projection, resulting in a single projected image. A binary mask of this image was created using the IsoData threshold method in ImageJ^80^. The outline of the probe was manually drawn onto the mask. The probe coverage percentage was calculated by determining the ratio of nonzero pixels representing the threshold GFP signal within the outline to the total number of enclosed pixels using ’Measure’ function.

### 2.6 Statistical analysis

Statistical analyses were performed using GraphPad Prism software (version 8.0.0 for Windows, GraphPad Software, San Diego, California, USA, www.graphpad.com). N-values represent individual animals, while n-values represent individual cells. Error bars are presented as means ± s.e.m. A significance level of α=0.05 was used for all analyses. To compare migration, T-index, D-index, surveillance, density, vessel diameter, and probe coverage between animal groups, two-way ANOVAs or mixed-effects models were employed, followed by Holm-Sidak’s or Tukey’s post hoc comparisons. Welch’s t-test was used to evaluate the effect of LIPUS on average vessel diameter on day 0 compared to control group. Bonferroni correction was applied where appropriate to correct for multiple comparison errors. Logistic regressions were applied to analyze ramification index data, while linear regressions were used to assess the spatial features of migration and surveillance.

## 3. Results

### 3.1 Experimental setup for LIPUS stimulation and two-photon imaging of microglia and vasculature

To optimize the application of LIPUS on brain tissue, we conducted *in vitro* testing of LIPUS parameters (Fig. 1a). The thermal effects of LIPUS were monitored via implantation of a thermocouple around 1.5 mm deep into the mouse cortex. A power supply voltage of 135 V for the transducer was carefully selected to ensure that the temperature change was below 1°C throughout the LIPUS treatment (Fig. 1b). In the LIPUS group (N=7), animals were treated with LIPUS followed by the two-photon imaging on day 0, 1, 2, 3, 4, 5, 6, 7, 14, 21, 28 post microelectrode implantation (Fig. 1c). Meanwhile, in the control group (N=7), animals underwent identical two-photon imaging sessions without LIPUS treatment. For LIPUS treatment, the transducer was positioned 14 mm from the cover glass with a PVA cone attached (Fig. 1d).

Preceding the insertion of the microelectrode, microglia exhibited a uniform distribution, and no discernible morphological alterations were observed in the cortical tissue caused by the craniotomy. However, the inherent limitations of two-photon microscopy have historically posed challenges when studying the tissue response to electrodes inserted perpendicularly into the brain. These challenges have been previously addressed in published work^22^. Specifically, a probe was inserted into the cortex at an angle of 30° and secured with a chronic imaging window (Fig. 1e) to enable two-photon visualization of fluorescently labeled microglia. To minimize bleeding, the insertion process was carefully conducted to avoid large surface vasculature, and sealing techniques were employed to preserve the region of interest during chronic imaging sessions^42,68,81^. The laser power was kept at ∼20 mW (never exceeding 40 mW) to prevent thermal damage. Photomultiplier tube (PMT) settings were adjusted accordingly for best image quality. For quantification purposes, a region of interest located 300 μm adjacent to the probe shank was selected (Fig. 1e). Furthermore, intraperitoneal administration of SR101 was performed to observe changes in the vasculature (magenta in Fig. 1e). These experimental setups confirmed that LIPUS could be safely delivered to the brain and allow for the chronic imaging of microglia activity (green in Fig. 1e) using two-photon microscopy.

### 3.2 LIPUS increased microglial migration velocity on day 1 and day 3

Microglial migration toward the injury site represents a key feature of the inflammatory response following microelectrode implantation^22,40,41^. Given the potential of LIPUS to attenuate the inflammatory response, our initial investigation focused on whether LIPUS could influence microglial migration. To explore this possibility, we aligned the images taken at different time points (see Methods 2.4). Subsequently, we generated a composite image in which the image from the earlier time point was rendered in magenta, while the image from the later time point was displayed in green. This alignment allowed us to estimate the microglial migration by measuring the displacement of each cell within the composite image during the interval between these two time points. The analysis revealed that LIPUS treatment significantly increased the microglial migration velocity on day 1 (1.35±0.07 vs 0.59±0.04 µm/hr) and day 3 (1.62±0.06 vs 0.96±0.05 µm/hr) but led to a reduction in velocity from day 4 to day 6 (Fig. 2b).

To visualize the spatial characteristics of microglia migration, the velocity of each individual microglia was plotted against the corresponding distance from the implantation site (Fig. 2c-f). Generally, there was a negative slope in the scatter plot, indicating that migration velocity tends to decrease as the distance from the implantation site increases. Notably, microglia close to the implantation site exhibited a higher velocity in the LIPUS group on day 1, as evidenced by a significantly higher intercept in the linear regression model (Fig. 2c). Our analysis demonstrated that LIPUS treatment significantly increased the velocity of microglia migration toward the implantation site during the early time points, but it resulted in a decrease in velocity at later stages (after day 4). These findings shed light on the directed movement of microglia toward the injury site and the benefits of LIPUS on boosting their migratory behavior.

### 3.3 LIPUS reduced the morphological activation of microglia within 100 µm from the microelectrode on day 6

After observing significant differences in microglial migration toward the injury site, we questioned whether LIPUS could also attenuate microglial morphological activation^22^ resulting from microelectrode implantation. We classified the microglia into two stages: transitional stage (0) and ramified stage (1) based on previous studies^22,40,45^. Microglia were sampled within a range of 0 to 300 µm from the probe and fitted into a logistic regression model (Fig. 3a). The Y-axis represents the percentage of ramified microglia at each distance bin (50: 0-50 µm, 100: 50-100 µm, 150: 100-150 µm, 200: 150-200 µm, 250: 200-250 µm, 300: 250-300 µm). Values closer to 1 indicates a higher proportion of ramified microglia, suggesting less morphological activation. Generally, ramification values increased with distance from the electrode surface in both LIPUS and control groups, confirming that morphological activation decreased with distance. To determine the suitability of fitting microglial ramification with a logistic regression model, the receiver operating characteristic (ROC) curve was plotted (Fig. 3b). The ROC curve was created by plotting the true positive rate against false positive rate for various threshold settings used to classify observations as positive or negative (detailed calculation documented here^72^). The diagonal random classifier line (black dashed line) indicates the performance expected from a classifier that makes predictions randomly, with no regard for the underlying data distribution. The closer distance between ROC curve and random classifier line Indicates a less suitability for fitting ramification data with a logistic regression model. Interestingly, the ROC curve began to approach the random classifier line (black dashed line) by day 6 in the LIPUS group and by day 7 in the control group (Fig. 3b), which could be due to the presence of more ramified microglia near the microelectrode starting day 6. The transition from a transitional state to a ramified state occurred earlier in the LIPUS group (on day 6) compared to the control group (on day 7), implying that LIPUS advanced the microglial transition from transitional/activated state to ramified state.

**Figure 3.**
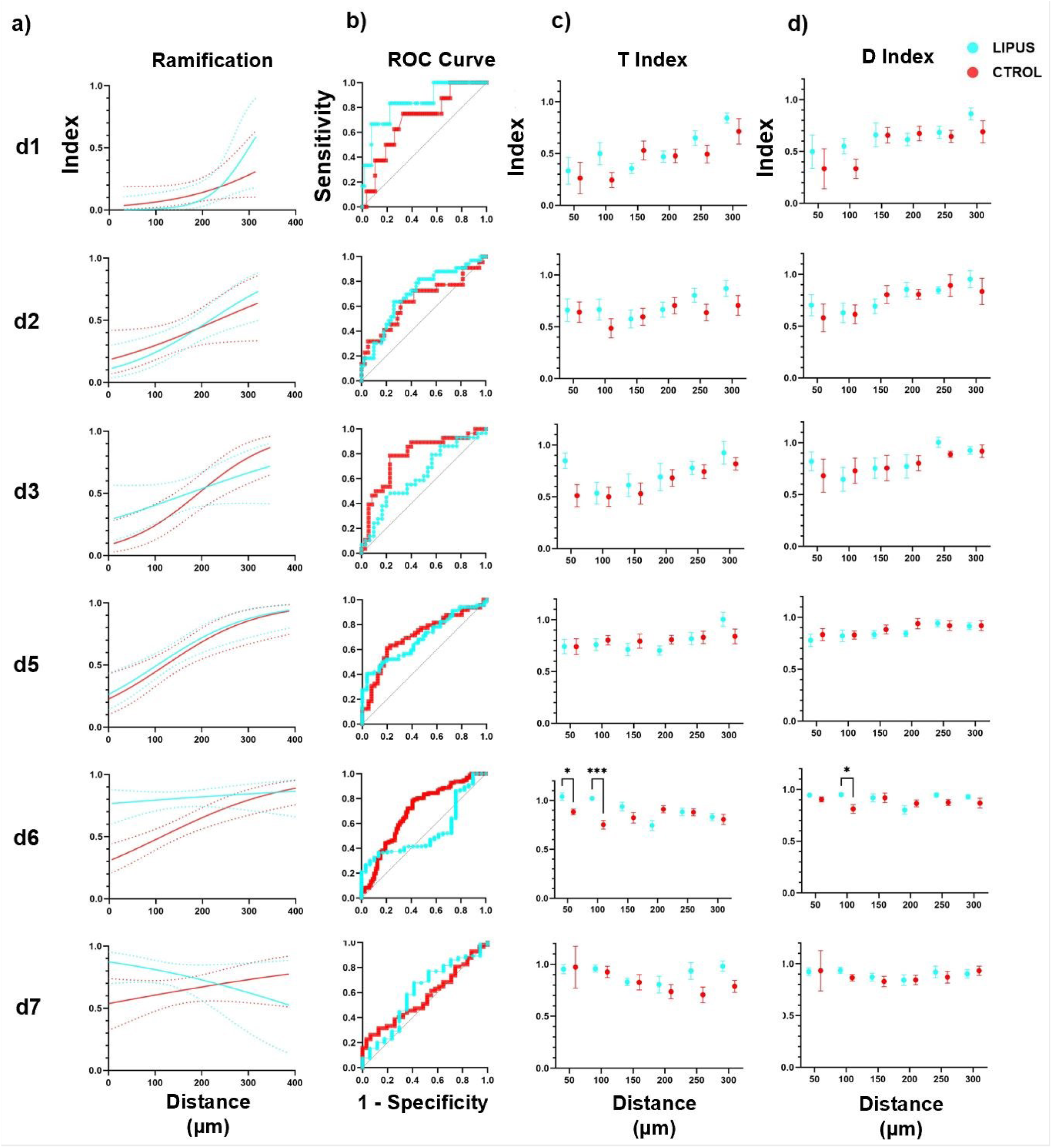
LIPUS attenuated microglial activation on day 6. **a)** Logistic regression analysis of microglia ramification over distances. The Y-axis represents the predicted percentage of ramified microglia using the logistic regression model. The dotted lines represent the 95% confidence bands. **b)** Receiver Operating Characteristic (ROC) curve for each logistic regression model used in panel **a)**. The proximity to the black dashed line indicates the model’s performance. The closer distance between ROC curve and the random classifier line (black dashed line) indicates a less suitability for fitting ramification data with a logistic regression model. Based on the ROC curves, the microglia ramification data was unsuitable for fitting into the logistic regression model on day 6 in the LIPUS group and on day 7 in the control group. **c)** T-index calculation, which assesses the degree of activation based on the length of the most prominent leading versus lagging microglia process relative to the electrode. Error bars indicate the standard errors. **d)** D-index calculation, which measures the degree of activation based on the number of leading versus lagging microglia processes relative to the electrode. Across spatial bins, the T-index and D-index generally decreased with time until day 3, indicating a higher degree of microglia activation. Starting from day 5, both indices began to increase. Notably, LIPUS treatment significantly increased both the T-index and D-index on day 6 (Šídák’s multiple comparisons test, * *p*<0.05, *** *p*<0.001), providing clear evidence of attenuated microglial morphological activation. Error bars indicate the standard errors.

**Figure 4.**
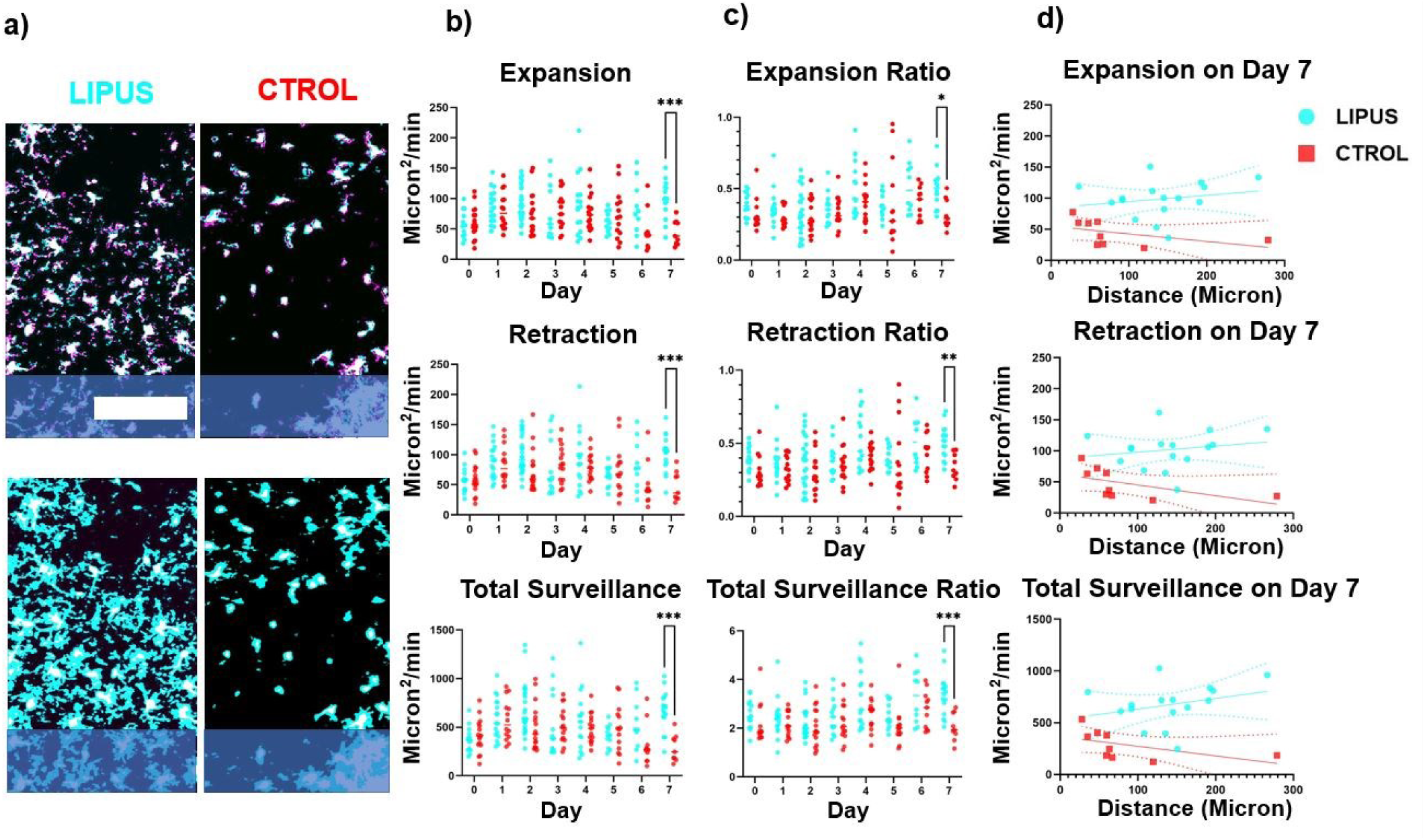
LIPUS increased the surveillance area expansion/retraction and total surveillance area of microglia on day 7. **a)** Top two rows: Examples of two-photon processed images showing microglia expansion (cyan), retraction (magenta), stable part (white) over a 1-minute interval. Bottom two rows: Examples of two-photon processed images displaying the total surveillance area (cyan) and stable part (white) of microglia over a 10-minute period. **b)** Quantification of expansion, retraction, and total surveillance area. LIPUS treatment significantly increased the expansion, retraction, and total surveillance area on day 7 (Šídák’s multiple comparisons test, *** *p*<0.001). **c)** The expansion, retraction, and surveillance normalized by the stable part of microglia. LIPUS treatment significantly increased the expansion ratio, retraction ratio, and total surveillance ratio on day 7 (Šídák’s multiple comparisons test, * *p*<0.05, ** *p*<0.01, *** *p*<0.001). **d)** Spatial characterization of microglia expansion/retraction and surveillance on day 7. Scale bar = 100 µm.

The degree of microglia activation was further evaluated using a transitional (T-) and directionality (D-) indices (see Methods 2.5.2), based on the length or number, respectively, of leading (towards) versus lagging (away) microglia processes, as described in previous studies^40^. Indices closer to 1 indicate a ramified state as there are equal length or number of processes facing toward and away from the probe, while indices closer to 0 suggest a more activated state due to a greater length or number of processes preferentially oriented towards the implant. Occasionally, there are values greater than one, indicating a preferred orientation of processes away from the probe, potentially due to population variability or directed process extension towards a damaged blood vessel caused by tissue strain or inflammation^82–84^. Smaller values for T-index and D-index indicated a higher level of microglial activation. Similarly high levels of microglia activation were seen near the implantation site in both LIPUS and control groups. This observation was consistent with previous findings^40,42^, indicating that the morphological activation of microglia was more pronounced in proximity to the implantation site (Fig. 3c-d). Notably, on day 6, the microglia processes exhibited significantly greater length and a higher T-index in the LIPUS group compared to the control group (Fig. 3c: within 50 µm, 1.04±0.04 vs 0.88±0.03, *p*<0.05; 50 – 100 µm, 1.02±0.02 vs 0.75±0.04, *p*<0.001). On the same day, the LIPUS group showed a significantly greater number of processes and a higher D-index compared to the control group (Fig. 3d: 50 – 100 µm, 0.95±0.01 vs 0.81±0.04, *p*<0.05). Therefore, our results revealed that LIPUS suppressed the morphological activation of microglia, especially on day 6 within 100 µm from the implantation site.

### 3.4 LIPUS increased the expansion/retraction and total surveillance of microglia on day 7

After observing significant LIPUS-induced differences in microglial migration and morphology, we asked whether those changes affect microglial surveillance of the surrounding tissue areas. To answer this question, we evaluated the continuous surveillance activity of microglia^85^. The average surveillance area expansion (blue in top row)/retraction (magenta in top row) and total surveillance area (blue in bottom row) were analyzed (Fig. 4a) based on previously published methods^73^. Specifically, the average expansion/retraction represents the rate of surveillance, while the total surveillance reflects the area monitored by microglial processes. The expansion/retraction speed and the total surveillance area initially increased then started to decrease from day 2 (Fig. 4b). Interestingly, LIPUS significantly increased the expansion (93.15±8.77 vs 44.50±6.86 µm^2^/min), retraction (101.84±7.58 vs 47.80±8.13 µm^2^/min), and total surveillance area (673.29±50.81 vs 286.43±46.21 µm^2^ per 10 mins) compared to control group on day 7 (*p*<0.001).

While we observed an increased surveillance activity of microglia in LIPUS group, we want to determine whether this was due to the larger soma area of microglia in the LIPUS group. It was noted that microglia near the implantation site were smaller compared to those away from the implantation site^86^. To account for this size effect, the average expansion/retraction speed and the total surveillance area were normalized by the size of the stable part of the microglia (Fig. 4c). A significant increase in the average expansion speed (49.64±3.24 vs 32.40±3.50%/min, *p*<0.05), retraction speed (51.01±2.82 vs 33.82±3.04%/min, *p*<0.01), and the total surveillance area (338.17±21.05 vs 205.59±19.02% per 10 mins, *p*<0.001) was still observed in the LIPUS group on day 7 (Fig. 4c). The spatial analysis of microglia expansion/retraction and surveillance indicated that LIPUS treatment increased the average expansion/retraction and total surveillance area on day 7 across all distances compared to the control group (Fig. 4d). Together, our findings demonstrate that LIPUS increased the microglia directed movements without affecting ramification or surveillance at earlier points (day 1, day 3), while promoting ramification and surveillance at later time points (day 6, day 7).

### 3.5 LIPUS reduced the probe coverage from day 6

We next calculated how LIPUS treatment affects microglial encapsulation of the microelectrodes as we have done previously with other interventions^22,40,42,45^. We quantified the percentage of probe coverage by calculating the ratio of the area occupied by GFP-positive microglia to the area of tissue directly above the probe shank (20 µm z-projection)^40,45^. A higher percentage of probe coverage indicates greater microglia encapsulation. We observed a rapid increase in the percentage of probe coverage in both groups from day 0 to day 5. However, the LIPUS group exhibited faster stabilization and reduced encapsulation of the probe around day 6 (Fig. 5b, 22.34±3.05 vs 39.08±4.41%, *p*<0.01), compared to the control group. By day 28, the LIPUS group had a significantly lower percentage of probe coverage compared to the control group (15±5.80% vs 51.68±2.58%, *p*<0.001). Taken together, LIPUS treatment leads to a faster stabilization (as early as day 6) and lower percentage of probe coverage by microglia, indicating reduced microglial encapsulation of microelectrodes.

**Figure 5.**
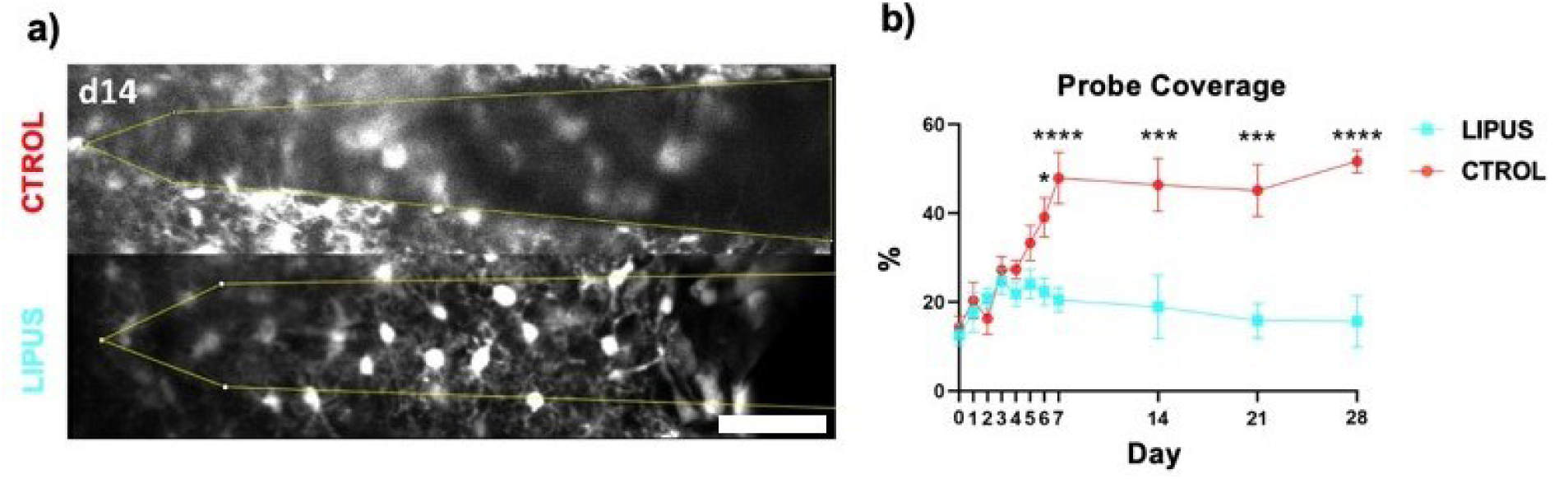
LIPUS attenuated microglial encapsulation of intracortical microelectrode starting day 6. **a)** Representative two-photon images on day 14 revealed that microglia extensively covered the probe in the control group, whereas in the LIPUS group, microglia displayed distinct processes with reduced overlap. **b)** The percentage of microglial surface coverage was quantified up to 20 µm above the surface of the implant (yellow outline in a), comparing LIPUS and control groups. LIPUS treatment resulted in a lower level of probe coverage, which occurred around day 6, and remained consistently lower than that in the control group (Šídák’s multiple comparisons test, * *p*<0.05, *** *p*<0.001, **** *p*<0.0001). Scale bar = 50 µm.

### 3.6 LIPUS reduced the number of vessel-associated microglia on day 7

The implantation of microelectrodes not only induces microglia encapsulation but also leads to the leakage of BBB^55^. This increased BBB leakage is temporally correlated with enhanced microglia-vessel interactions^25^. To assess the impact of LIPUS on microglia-vessel interactions, we quantified the total microglia population, vessel-associated microglia (defined as microglia extending at least one process onto vasculature), and their ratio over time post-implantation (Fig. 6). In both the LIPUS and the control group, the density of the total microglia and vessel-associated microglia decreased from day 0 to day 3, followed by an increase from day 3 to day 7 (Fig. 6b and Fig. 6c). Additionally, the vessel-associated microglia ratio peaked at 80% by day 2 and then gradually decreased to around 40% on day 6 for both the LIPUS and the control group (Fig. 6d). However, while the LIPUS group maintained a vessel-associated microglia ratio of approximately 40% on day 7, the median ratio in the control group rose above 70% (Fig. 6d, 40.43±3.87 vs 70.67±6.15%, *p*<0.0002). To account for individual differences, we analyzed the ratio change by subtracting the ratio on day 0. Notably, LIPUS treatment led to a significant decrease in the ratio change of vessel-associated microglia on day 7 (Fig. 6e, -32.62±6.18 vs 3.72±4.07%, *p*<0.0001). These findings suggest that LIPUS may mitigate the accumulation of microglia on nearby blood vessels during the transition from the acute phase to the chronic phase.

### 3.7 LIPUS reduced the diameters of cerebral blood vessels on day 28

While the role of microglia in the regulation of vasculature has only recently begun to be elucidated^25,87^, it is evident that the interaction between microglia and the vasculature does not occur randomly^87^. We have observed in our previous studies that cerebral blood vessels dilate following microelectrode implantation^39,41,49,81^, which was temporally correlated with the activation of microglia. Hence, we wondered whether the reduction in microglial activation and vessel-associated microglia ratio by LIPUS might contribute to the prevention of blood vessel dilation chronically. The diameters of blood vessels near the implantation site were evaluated using VasoMetrics ImageJ plugin^77^ (Fig. 7). Similar to previous studies^39,41,49,81^, we found a steady increase in the average blood vessel diameter during the first 6 days. From day 7 to day 28, the control group exhibited a slight increase, while the LIPUS group showed a slight decrease (Fig. 7b). Interestingly, LIPUS significantly reduced the vessel diameter on day 28 (Fig. 7b, 3.95±0.13 vs 4.76±0.14 µm, *p*<0.05). To eliminate the possibility that the reduction in blood vessel diameter was a result of the initial vessel selection, we carefully selected blood vessels with similar diameters on day 0 in both groups (Fig. 7c). LIPUS significantly reduced vessel diameter only after LIPUS stimulation (Fig. 7c, 4.37±0.17 vs 4.54±0.16 µm, *p*<0.05). Since LIPUS induced a reduction in vessel diameters, we want to know if LIPUS also affected the density and structure of the vessels. We did not detect a significant difference in vessel area coverage (Fig. 7d), indicating the density of vessels was unaffected by LIPUS. In addition, tortuosity (Fig. 7e), vessel branch length (Fig. 7f and g) and vessel branch number (Fig. 7h) quantified by skeletonize ImageJ macro^77,78^ were not significantly different between LIPUS and control group, indicating vessel structures were unaffected. Taken together, our findings suggest that LIPUS treatment reduces vessel dilation but does not affect the structure of the cerebral blood vessels.

**Figure 6.**
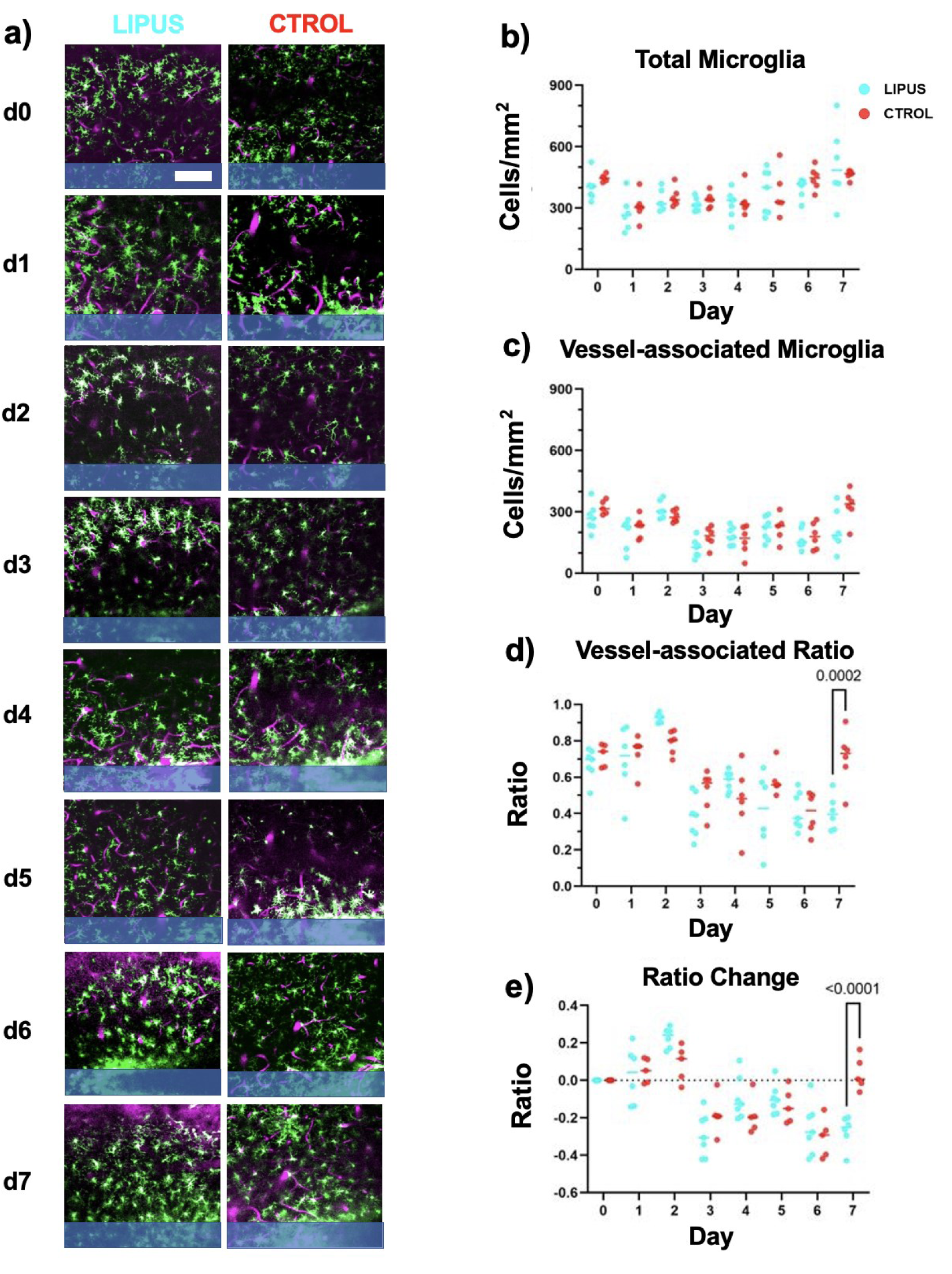
LIPUS reduced the number of vessel-associated microglia on day 7. To identify vessel-associated microglia, a 22-mm Z stack was analyzed to identify microglial processes that exhibited at least one attachment to the blood vessel. **a)** Representative two-photon images showing microglia (green) and vasculature (magenta) from day 0 to day 7. **b)** The total microglia density (including non-vessel-associated and vessel-associated microglia) initially decreased up to day 3 and then returned to the same level as day 0 in both LIPUS and control groups. **c)** Vessel-associated microglia density also exhibited a similar trend, decreasing up to day 3 and then returning to the baseline level. **d)** The vessel-associated ratio was calculated by dividing the number of vessel-associated microglia by the total number of microglia. LIPUS treatment appeared to increase the vessel-associated microglia ratio on day 2 and significantly decrease it on day 7 (Šídák’s multiple comparisons test). **e)** To account for individual variability, the ratio change was calculated by subtracting the vessel-associated ratio on each day from the vessel-associated ratio on day 0 for each animal. LIPUS treatment significantly reduced the vessel-associated microglia ratio on day 7 (Šídák’s multiple comparisons test). Scale bar = 100 µm.

**Figure 7.**
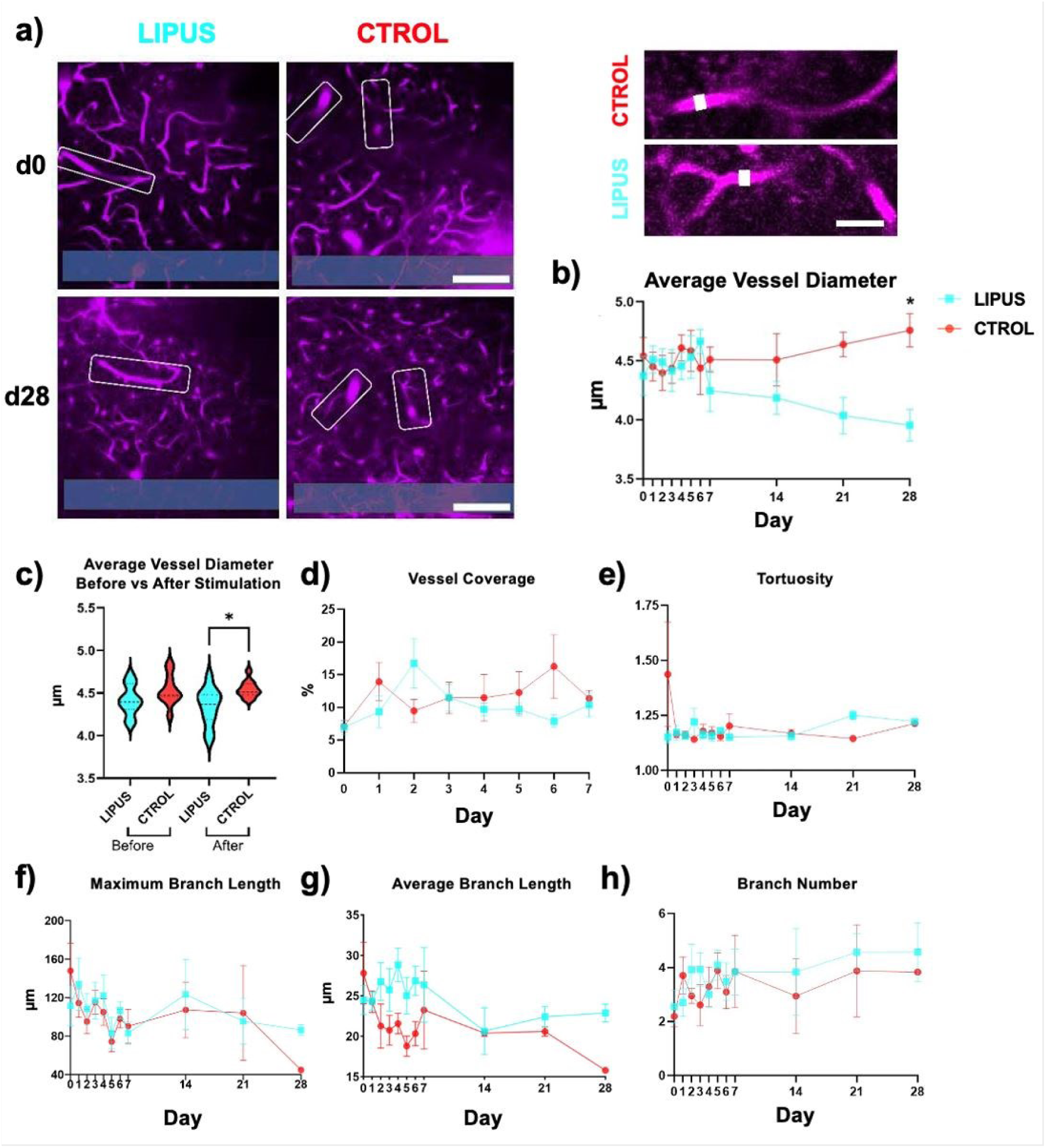
LIPUS affected morphology of the vasculature after chronic probe insertion. **a)** Left images: two-photon microscopy images of LIPUS and control group on day 0 and day 28. White boxes denote the analyzed vasculatures. The probe is outlined in blue at the bottom of the images. Scale bar = 100 µm. Right images: magnification of blood vessels on day 28 in both groups. Scale bar = 20 µm. **b)** Average blood vessel diameter over 28 days. LIPUS treatment slightly decreases the average vessel diameter on day 28 (Tukey’s multiple comparison test, * *p*<0.05). **c)** Average vessel diameter before and after stimulation on day 0. Significant reduction in average blood vessel diameter in the LIPUS group was observed only after stimulation compared to the control group (Welch’s t test, * *p*<0.05). **d)** No significant difference between groups was detected in vessel area coverage percentage. **e)** Tortuosity measures the level of twisting or distortion of the vessels. No significant difference between groups was detected in tortuosity. **f)** Maximum blood vessel branch length showed no significant difference between groups. **g)** A higher average branch length was observed in LIPUS group compared to control group from day 2 to day 7 and day 28, however, no significant difference was detected. **h)** No significant difference between groups was detected in number of blood vessel branches per vessel.

## Discussion

LIPUS has been shown to be a promising tool for suppressing inflammation^88^ and microglia activation^89^. Our findings indicated that LIPUS treatment resulted in increased microglia migration on day 1 and day 3 (Fig. 2), more ramified morphology on day 6 (Fig. 3), enhanced microglia surveillance on day 7 (Fig. 4), decreased probe coverage from day 7 (Fig. 5), reduced vessel-associated microglia ratios on day 7 (Fig. 6), and attenuated dilation of blood vessels on day 28 (Fig. 7).

Microglia-mediated neuroinflammation has been hypothesized to impede tissue-microelectrode integration. The BBB is compromised during microelectrode implantation, resulting in infiltration of blood contents which causes damage to neural tissue^49^. Molecules, such as fibrin^90^ and ATP^91^, released from the damaged area attract microglia, prompting them to migrate toward the injury site^49^. Previous studies showed that microglia initiate their migration within 6 – 24 hours post implantation^41,43–45^. Interestingly, our results show that microglia migration speed is higher in the LIPUS group compared to control group on day 1 and day 3 (Fig. 2). The faster migration of microglia toward the implantation site suggests that LIPUS could promote microglia to facilitate the closure of the wounds caused by microelectrode implantation, thereby potentially mitigating further neuronal damage by reducing the infiltration of blood contents^92^. If the injury and inflammation are not resolved timely, events such as oxidative stress^93^ and microglia priming^94^ can lead to persistent activation of microglia. As a result, activated microglia secrete proinflammatory cytokines that lead to the formation of a glial scar^51^. The glial scar not only displaces neurons from the recording range^51,95^ but also forms a physical barrier, inhibiting ion diffusion and preventing the propagation and detection of neural signals^37,49,68,96,97^. Our results indicate that enhanced microglia repair via LIPUS at acute timepoints (day 1 and day 3) (Fig. 2) limit propagation of injury, thereby reducing microglia activation at chronic time points (day 6) (Fig. 3). The decrease in microglia activation at chronic stages may reduce the release of proinflammatory cytokines^64^ and NO^98^, while also mitigating harmful events such as lysosomal dysfunction^99,100^, peroxidated lipid accumulation^101^, and ferroptosis^102^. These proinflammatory processes may coordinate with over proliferation of microglia, leading to microgliosis and increased microglial probe coverage as the chronic neuroinflammation progresses^103–105^. In line with the reduced microglial activation, we observed reduced probe coverage at later stages (starting day 6) in LIPUS group (Fig. 5), indicating better interfacing between tissue and implanted microelectrode^45^.

Previous studies have highlighted the dilation of blood vessels post implantation^39,41^, potentially due to the release of NO^98,106^. This abnormal vasodilation has been linked to exacerbated brain swelling^107^, indicating a disruption in neurovascular coupling. An earlier study demonstrated that LIPUS promoted vascular remodeling and increased the vessel diameter and length on day 3 following distal middle cerebral artery occlusion^108^. Even though the vessel length was consistently higher in our LIPUS group from day 2 to day 6 (Fig. 7g), we did not detect a significant difference between groups on each individual day (with the lowest *p*-value on day 4: *p*=0.24), neither in vessel length nor in vessel diameter. The chronic reduction in vessel diameter on day 28, rather than an acute change (Fig. 7b), implies that the benefits of LIPUS treatment are less likely to result from vascular remodeling. Instead, the accelerated microglia repair (Fig. 2) and reduction in persistent microglia activation (Fig. 4) may contribute to the long-term reduction in NO release^28^, preventing the dilation of blood vessels. The observed decline in vasodilation at a chronic time point (Fig. 7b) suggests that LIPUS treatment may help preserve neurovascular coupling, which is essential to provide energy for neuronal firing^109^. Future studies that record neural signals after LIPUS treatment could elucidate whether improved tissue-microelectrode integration enhances device recording performances.

Neuroinflammation involves several signaling pathways that can be regulated by mechanosensitive and thermosensitive channels^110,111^. Ultrasonic wave delivered by LIPUS can open these channels, thereby mediating neuroinflammation. Whether the effects of LIPUS could be beneficial or detrimental to resolution of neuroinflammation largely depends on the underlying mechanisms and pathways it engages. For example, the activation of TRPA1 receptors may exacerbate myelin damage and reduce cognitive outcomes^112^, while activating TRPV1 receptor facilitates myelin repair which are important mechanisms for functional recovery^113^. Several channels and receptors can modulate microglial activity including microglial migration, ramification, and surveillance^85,113–115^. Notably, the activation of TRPV1 receptors boosts the microglia migration^113^, potentially contributing to the observed increase in microglial migration on day 1 and day 3 (Fig. 2). Similar to TRP channels, the activation of mechanosensitive channels such as Piezo1^114^ can also result in enhanced microglial migration (Fig. 2) via cytoskeleton remodeling^116–120^.

Furthermore, LIPUS may amplify the release of ATP^121^, possibly originating from the injury site^122^, connexin 43 hemichannels in astrocytes^123^ and Pannexin 1 channels in vasculature^87^. ATP, in turn, activates P2Y12 receptors on microglia, thereby increasing the speed of their directed movements toward damaged area^115^. Interestingly, we did not detect significant increase in microglial ramifications (Fig. 3) and surveillance (Fig. 4) on day 1 and day 3, even with enhanced migration speed of microglia (Fig. 2) in the LIPUS group. The ramification and surveillance of microglia are primarily regulated by one of the two-pore domain potassium (K2P) channels – THIK-1^85^. Since THIK-1 can be potentiated by activation of P2Y12 receptors^85^, an increase in ramification and surveillance would typically coincide with any ATP-induced enhancement in microglial migration speed. The fact that we observed only increased migration (Fig. 2), but no significant improvement in ramification (Fig. 3) and surveillance (Fig. 4), suggests that LIPUS was less likely to influence microglial activity through ATP release, as opposed to the activation of Piezo1 or TRPV1 during the acute phase.

Moreover, microelectrode implantation leads to the infiltration of blood contents such as fibrinogen, due to the rupture of BBB^49^. The immunoreactivity of fibrinogen near the microelectrode was significantly higher than the distal region around 1 week^124^. Previous studies have shown that fibrinogen can trigger microglial activation^125,126^, leading to generation of reactive oxidative stress in microglia^90^. The elevated oxidative stress accompanied by dysfunctions in microglia metabolism can lead to impaired microglial clearance ability^127^. While the overproliferation of microglia may transiently compensate for the impaired clearance capabilities, it ultimately contributes to microglial senescence^128^. These underlying processes could explain the accumulation of activated microglia near the microelectrode, particularly at early stages (before day 6) (Fig. 4). As the fibrin or cell debris was cleared, the microglia near the microelectrode started to transition back to a ramified state (Fig. 4).

Interestingly, this transition happens earlier in the LIPUS (starting day 6) compared to control group. Consequently, on day 6 (Fig. 4c), we observed that microglia in the LIPUS group exhibited longer process lengths and a higher number of processes. Mechanistically, LIPUS can facilitate this transition by potentially breaking down fibrin polymers^129^, thereby reducing the burden of clearance, or by enhancing glycolytic metabolism^127^ through the activation of mechanosensitive channels^130^. Future experiments to apply LIPUS after scar tissue formation may help test these hypotheses and explore the potential to mitigate the scar formation during the chronic phase following microelectrode implantation. In summary, our data on microglial migration (Fig. 2), ramification (Fig. 3), and surveillance (Fig. 4) suggest that LIPUS may benefit resolution of microglia-mediated neuroinflammation via activation of mechanosensitive channels during the acute phase, promoting fibrin thrombolysis and boosting microglia metabolism at later stages.

LIPUS not only has the potential to improve tissue-microelectrode bio-integration for BCIs, but also provide a new opportunity for treating other dysfunctions in biological processes^131,132^, CNS injuries^133^ and neurodegenerative diseases^134–136^. For example, in response to ischemic injury, microglia migrate toward the injury site to phagocytose cell debris and protect damaged cells, reducing the inflammation caused by dead cell corpses^137^. However, ischemia could lead to oxygen and glucose deprivation which reduces the microglia migration speed^138^, adversely impacting microglia clearance and injury recovery. Increased microglia migration by LIPUS (Fig. 2) not only accelerates this clearance process but also provides better protection for neurons damaged by oxygen-glucose deprivation^139^. Furthermore, surveillance of microglia may be impaired in ischemia^138^ as reduced blood flow can limit the expansion and retraction of microglia processes^140^. LIPUS intervention, which increases expansion and retraction (Fig. 4), potentially enhances immune surveillance capabilities^19^. The enhancement of surveillance (Fig. 4) could promote microglia to sense subtle changes in the environment, collect tissue debris, and phagocytose leaked blood component^141–143^.

Moreover, in the context of Alzheimer’s disease, it has been shown that vessel-associated microglia constitute 60% of the total microglia population within the AD-affected brains^144^. This percentage is notably higher than the 30% typically found in healthy brain^25^. A decrease in the presence of vessel-associated microglia has been linked to lower levels of amyloid-beta (Aβ) and attenuated cognitive impairment^144^. Future experiments are necessary to investigate whether the increased presence of vessel-associated microglia shares a common cause between AD-affected brains and brains with implanted microelectrodes. Nevertheless, the reduction in vessel-associated microglia suggests that LIPUS treatment has the potential to improve cognitive performance in AD patients. Additionally, fewer vessel-associated microglia in a later stage could prevent the disruption of the BBB^25^, reducing the neuroinflammation induced by α-synuclein in Parkinson’s disease^145^. Furthermore, microglia in aged individuals tend to exhibit de-ramification, a process linked to aging^146^. From our results, microglia in LIPUS group display a greater number of processes, and these processes are longer, exhibiting a more ramified morphology (Fig. 3). Together with recent literature demonstrating that LIPUS can reduce senescent marker (P16^INK4a^) in aged mouse^131^, LIPUS treatment appears to have potential anti-aging effects on microglia.

To optimize the application of ultrasound stimulation for both the bio-integration of BCIs and the treatment of neurodegenerative diseases, future studies should explore the parameter space of ultrasound stimulation to achieve optimal therapeutic benefits. These parameters include intensity^147^, frequency^148^, pulse repetition frequency^149^, and duty cycle^150^. Among these parameters, it was observed that higher intensity ultrasound led to an increase in blood flow^151^. However, excessively high intensity could potentially breach the BBB^152^. Furthermore, different frequencies of ultrasound have the potential to induce distinct biological responses. For example, lower frequencies (e.g. 350 kHz) activate TRPA1 channels, while higher frequencies (43 MHz) stimulate piezo1 channels^148^. Moreover, the adjustment of pulse repetition has the potential to selectively modulate excitatory and inhibitory neurons^149^, providing a promising approach to precise neuromodulation. However, it is important to consider that an increase in duty cycle could result in a linear increase in temperature^150^. Therefore, careful consideration is necessary to ensure optimal therapeutic benefits while avoiding excessive temperature rises during ultrasound stimulation. In addition to parameter considerations, the variations in bone density within the skull pose challenges for precise targeting of LIPUS to the intended treatment area^153^. Future studies should focus on refining the models of ultrasound propagation through the skull^154^ and developing phase arrays^155^ to steer focal ultrasound to improve the precision of LIPUS treatment.

## Conclusion

This study investigated the potential of low-intensity pulsed ultrasound (LIPUS) as a therapeutic intervention for mitigating neuroinflammation induced by the implantation of intracortical microelectrodes. We found that LIPUS treatment effectively promotes wound closure by enhancing microglial migration toward the implantation site on day 1 and day 3. While the microglial ramification and surveillance remained unchanged until day 5, LIPUS treatment significantly reduced the microglial activation on day 6. Furthermore, microglial expansion/retraction area and the total surveillance area were improved in the LIPUS group on day 7. Additionally, LIPUS significantly reduced the microglial encapsulation of the microelectrode starting day 6, lowered vessel-associated microglia ratio on day 7, and reduced the average vessel diameter on day 28. These findings suggest that LIPUS treatment not only has the potential to attenuate microglial encapsulation of intracortical microelectrodes but also boost the healing process and mitigate neuroinflammation in other forms of brain injury.

## Supplementary tables

**Statistics for Figure 2.**

| Day | Animal (N) | Cell (n) | *p* value |
| --- | --- | --- | --- |
| 1 | LIPUS: 6, CONTROL: 5 | LIPUS: 36, CONTROL: 29 | $p<0.0001$ |
| 3 | LIPUS: 7, CONTROL: 4 | LIPUS: 40, CONTROL: 23 | $p<0.0001$ |
| 4 | LIPUS: 7, CONTROL: 4 | LIPUS: 38, CONTROL: 22 | $p<0.0001$ |
| 5 | LIPUS: 7, CONTROL: 4 | LIPUS: 37, CONTROL: 23 | $p<0.0001$ |
| 6 | LIPUS: 5, CONTROL: 4 | LIPUS: 23, CONTROL: 22 | $p<0.05$ |

**Statistics for Figure 3.** T-index on Day 6.

| Distance | Animal (N) | Cell (n) | *p* value |
| --- | --- | --- | --- |
| within 50 μm | LIPUS: 5, CONTROL: 4 | LIPUS: 21, CONTROL: 40 | $p<0.05$ |
| 50 – 100 μm | LIPUS: 5, CONTROL: 4 | LIPUS: 29, CONTROL: 29 | $p<0.001$ |

**Statistics for Figure 3.** D-index on Day 6.

| Distance | Animal (N) | Cell (n) | *p* value |
| --- | --- | --- | --- |
| 50 – 100 μm | LIPUS: 5, CONTROL: 4 | LIPUS: 29, CONTROL: 29 | $p<0.05$ |

**Statistics for Figure 4.** on Day 7.

| Measurements | Animal (N) | Cell (n) | *p* value |
| --- | --- | --- | --- |
| Expansion | LIPUS: 4, CONTROL: 2 | LIPUS: 16, CONTROL: 9 | $p<0.001$ |
| Retraction | LIPUS: 4, CONTROL: 2 | LIPUS: 16, CONTROL: 9 | $p<0.001$ |
| Surveillance | LIPUS: 4, CONTROL: 2 | LIPUS: 16, CONTROL: 9 | $p<0.001$ |
| Expansion Ratio | LIPUS: 4, CONTROL: 2 | LIPUS: 16, CONTROL: 9 | $p<0.05$ |
| Retraction Ratio | LIPUS: 4, CONTROL: 2 | LIPUS: 16, CONTROL: 9 | $p<0.01$ |
| Surveillance Ratio | LIPUS: 4, CONTROL: 2 | LIPUS: 16, CONTROL: 9 | $p<0.001$ |

**Statistics for Figure 5.**

| Measurements | Day | Animal (N) | *P* value |
| --- | --- | --- | --- |
| Probe coverage | 6 | LIPUS: 7, CONTROL: 6 | $p<0.05$ |
| Probe coverage | 7 | LIPUS: 7, CONTROL: 6 | $p<0.0001$ |
| Probe coverage | 14 | LIPUS: 4, CONTROL: 6 | $p<0.0001$ |
| Probe coverage | 21 | LIPUS: 3, CONTROL: 5 | $p<0.0001$ |
| Probe coverage | 28 | LIPUS: 3, CONTROL: 4 | $p<0.0001$ |

**Statistics for Figure 6.**

| Measurements | Day | Animal (N) | *p* value |
| --- | --- | --- | --- |
| Vessel-associated ratio | 7 | LIPUS: 6, CONTROL: 6 | $p<0.0002$ |
| Ratio change | 7 | LIPUS: 6, CONTROL: 6 | $p<0.0001$ |

**Statistics for Figure 7.**

| Measurements | Animal (N) | Vessel (n) | *p* value |
| --- | --- | --- | --- |
| Average blood vessel diameter on day 28 | LIPUS: 3, CONTROL: 4 | LIPUS: 7, CONTROL: 5 | $p<0.0001$ |

## Acknowledgements

This work was supported by NIH R44MH131514, NIH R21EB028055, NIH R01NS094396, NIH R01NS105691, NIH R01NS129632, NIH R01NS115707 and a Diversity Supplement to this parent grant, as well as NSF CAREER 1943906 and an NSF GRFP. The authors would like to acknowledge valuable contributions and feedback on the manuscript from Stieger K, Chen K, Forrest A, Hughes C, Suematsu N, and Zhang G.

## Author information

### Contributions

Conceptualization: Tirko NN, Mulvihill M, Bagwell R, Kozai TDY.

Supervision: Tirko NN, Greaser J, Alsubhi AS, Gheres KW, Kozai TDY.

Methodology: Li F, Tirko NN, Gallego J, Patel R, Shaker E, Valkenburg GEV, Greaser J, Alsubhi AS, Gheres KW, Wellman S, Kozai TDY.

Investigation: Li F, Gallego J, Shaker E, Bashe D, Tirko NN, Alsubhi AS, Gheres KW, Kozai TDY.

Formal analysis: Li F, Gallego J, Patel R, Shaker E, Valkenburg GEV, Bashe D, Singh V, Gheres KW, Garcia Padill C.

Resources: Tirko NN, Greaser J, Gheres KW, Bagwell R, Mulvihill M, Kozai TDY.

Data Curation: Li F, Gallego J, Bashe D, Patel R, Shaker E, Valkenburg GEV, Singh V. Visualization: Li F, Gallego J, Patel R, Valkenburg GEV, Tirko NN, Gheres KW.

Writing – Original Draft: Li F.

Writing – Review & Editing: Li F, Gallego J, Shaker E, Singh V, Garcia Padill C, Wellman S, Tirko NN, Gheres KW, Kozai TDY.

Project administration: Li F.

Funding acquisition: Tirko NN, Mulvihill M, Bagwell R, Kozai TDY. <u>Corresponding authors</u>

Correspondence to Kozai TDY

## Ethics declarations

Tirko NN, Greaser J, Gheres KW, Bagwell R, Mulvihill M have financial stakes in Actuated Medical. However, it is important to note that the competing interests did not affect the design, methodology, or interpretation of the results in any way. We declare this competing interest to maintain transparency and to provide readers with full disclosure.

## Data availability

The data that support the findings of this study are available from the corresponding author upon reasonable request.

## References

1 Medzhitov, R. Origin and physiological roles of inflammation. Nature 454, 428–435 (2008).

2 Freire, M. O. & Van Dyke, T. E. Natural resolution of inflammation. Periodontology 2000 63, 149–164 (2013).

3 Grammas, P. Neurovascular dysfunction, inflammation and endothelial activation: implications for the pathogenesis of Alzheimer’s disease. Journal of neuroinflammation 8, 1–12 (2011).

4 Tarantini, S., Tran, C. H. T., Gordon, G. R., Ungvari, Z. & Csiszar, A. Impaired neurovascular coupling in aging and Alzheimer’s disease: contribution of astrocyte dysfunction and endothelial impairment to cognitive decline. Experimental gerontology 94, 52–58 (2017).

5 Petty, M. A. & Lo, E. H. Junctional complexes of the blood–brain barrier: permeability changes in neuroinflammation. Progress in neurobiology 68, 311–323 (2002).

6 McLarnon, J. G. A leaky blood–brain barrier to fibrinogen contributes to oxidative damage in Alzheimer’s disease. Antioxidants 11, 102 (2021).

7 Ownby, R. L. Neuroinflammation and cognitive aging. Current psychiatry reports 12, 39–45 (2010).

8 Ji, R.-R., Xu, Z.-Z. & Gao, Y.-J. Emerging targets in neuroinflammation-driven chronic pain. Nature reviews Drug discovery 13, 533–548 (2014).

9 Alghamri, M. S. et al. Targeting neuroinflammation in brain cancer: uncovering mechanisms, pharmacological targets, and neuropharmaceutical developments. Frontiers in Pharmacology 12, 680021 (2021).

10 Jayaraj, R. L., Azimullah, S., Beiram, R., Jalal, F. Y. & Rosenberg, G. A. Neuroinflammation: friend and foe for ischemic stroke. Journal of neuroinflammation 16, 1–24 (2019).

11 Kumar, A. & Loane, D. J. Neuroinflammation after traumatic brain injury: opportunities for therapeutic intervention. Brain, behavior, and immunity 26, 1191–1201 (2012).

12 Heneka, M. T. et al. Neuroinflammation in Alzheimer’s disease. The Lancet Neurology 14, 388–405 (2015).

13 Bjelobaba, I., Savic, D. & Lavrnja, I. Multiple sclerosis and neuroinflammation: the overview of current and prospective therapies. Current pharmaceutical design 23, 693–730 (2017).

14 Hirsch, E. C., Vyas, S. & Hunot, S. Neuroinflammation in Parkinson’s disease. Parkinsonism & related disorders 18, S210–S212 (2012).

15 Rayasam, A., Fukuzaki, Y. & Vexler, Z. S. Microglia-leucocyte axis in cerebral ischaemia and inflammation in the developing brain. Acta Physiologica 233, e13674 (2021).

16 Wu, F. et al. CXCR2 is essential for cerebral endothelial activation and leukocyte recruitment during neuroinflammation. Journal of neuroinflammation 12, 1–15 (2015).

17 Yang, Q. q. & Zhou, J. w. Neuroinflammation in the central nervous system: Symphony of glial cells. Glia 67, 1017–1035 (2019).

18 Li, Q. & Barres, B. A. Microglia and macrophages in brain homeostasis and disease. Nature Reviews Immunology 18, 225–242 (2018).

19 Bernier, L.-P. et al. Nanoscale surveillance of the brain by microglia via cAMP-regulated filopodia. Cell reports 27, 2895–2908. e2894 (2019).

20 Wahedi, U., Baba, M. Z., Praveen, T. K. & Subramanian, G. Damage-associated molecular patterns & m2 microglial polarization in Parkinson’s disease-a brief review. Journal of Medical Pharmaceutical and allied Science 11, 5289–5297 (2022).

21 Li, L. Z. et al. Potential microglia-based interventions for stroke. CNS Neuroscience & Therapeutics 26, 288–296 (2020).

22 Kozai, T. D. Y., Vazquez, A. L., Weaver, C. L., Kim, S.-G. & Cui, X. T. In vivo two-photon microscopy reveals immediate microglial reaction to implantation of microelectrode through extension of processes. Journal of neural engineering 9, 066001 (2012).

23 Brown, G. C. Neuronal loss after stroke due to microglial phagocytosis of stressed neurons. International journal of molecular sciences 22, 13442 (2021).

24 Chen, W. et al. Microglial phagocytosis and regulatory mechanisms after stroke. Journal of Cerebral Blood Flow & Metabolism 42, 1579–1596 (2022).

25 Haruwaka, K. et al. Dual microglia effects on blood brain barrier permeability induced by systemic inflammation. Nature communications 10, 5816 (2019).

26 Dudvarski Stankovic, N., Teodorczyk, M., Ploen, R., Zipp, F. & Schmidt, M. H. Microglia–blood vessel interactions: a double-edged sword in brain pathologies. Acta neuropathologica 131, 347–363 (2016).

27 Li, T. et al. Proliferation of parenchymal microglia is the main source of microgliosis after ischaemic stroke. Brain 136, 3578–3588 (2013).

28 Goodwin, J. L., Uemura, E. & Cunnick, J. E. Microglial release of nitric oxide by the synergistic action of β-amyloid and IFN-γ. Brain research 692, 207–214 (1995).

29 Van Mil, A. H. et al. Nitric oxide mediates hypoxia-induced cerebral vasodilation in humans. Journal of applied physiology 92, 962–966 (2002).

30 Butler, C. A. et al. Microglial phagocytosis of neurons in neurodegeneration, and its regulation. Journal of neurochemistry 158, 621–639 (2021).

31 Wilton, D. K. et al. Microglia and complement mediate early corticostriatal synapse loss and cognitive dysfunction in Huntington’s disease. Nature Medicine, 1–19 (2023).

32 Meneghetti, N. et al. Narrow and broad gamma bands process complementary visual information in mouse primary visual cortex. Eneuro (2021).

33 Mizuseki, K., Royer, S., Diba, K. & Buzsáki, G. Activity dynamics and behavioral correlates of CA3 and CA1 hippocampal pyramidal neurons. Hippocampus 22, 1659–1680 (2012).

34 Huber, D. et al. Multiple dynamic representations in the motor cortex during sensorimotor learning. Nature 484, 473–478 (2012).

35 Jia, X., Smith, M. A. & Kohn, A. Stimulus selectivity and spatial coherence of gamma components of the local field potential. Journal of Neuroscience 31, 9390–9403 (2011).

36 Flesher, S. N. et al. A brain-computer interface that evokes tactile sensations improves robotic arm control. Science 372, 831–836 (2021).

37 Collinger, J. L. et al. High-performance neuroprosthetic control by an individual with tetraplegia. Lancet 381, 557–564, doi:10.1016/S0140-6736(12)61816-9 (2013).

38 Hochberg, L. R. et al. Reach and grasp by people with tetraplegia using a neurally controlled robotic arm. Nature 485, 372–375, doi:10.1038/nature11076 (2012).

39 Savya, S. P. et al. In vivo spatiotemporal dynamics of astrocyte reactivity following neural electrode implantation. Biomaterials 289, 121784 (2022).

40 Dubaniewicz, M. et al. Inhibition of Na+/H+ exchanger modulates microglial activation and scar formation following microelectrode implantation. Journal of neural engineering 18, 045001 (2021).

41 Wellman, S. M. & Kozai, T. D. In vivo spatiotemporal dynamics of NG2 glia activity caused by neural electrode implantation. Biomaterials 164, 121–133 (2018).

42 Kozai, T. D., Jaquins-Gerstl, A. S., Vazquez, A. L., Michael, A. C. & Cui, X. T. Dexamethasone retrodialysis attenuates microglial response to implanted probes in vivo. Biomaterials 87, 157–169 (2016).

43 Sharon, A., Jankowski, M. M., Shmoel, N., Erez, H. & Spira, M. E. Inflammatory foreign body response induced by neuro-implants in rat cortices depleted of resident microglia by a CSF1R inhibitor and its implications. Frontiers in neuroscience 15, 646914 (2021).

44 Kozai, T. D. et al. Chronic tissue response to carboxymethyl cellulose based dissolvable insertion needle for ultra-small neural probes. Biomaterials 35, 9255–9268 (2014).

45 Eles, J. R. et al. Neuroadhesive L1 coating attenuates acute microglial attachment to neural electrodes as revealed by live two-photon microscopy. Biomaterials 113, 279–292 (2017).

46 Krukiewicz, K. Electrochemical impedance spectroscopy as a versatile tool for the characterization of neural tissue: A mini review. Electrochemistry Communications 116, 106742 (2020).

47 Frampton, J. P., Hynd, M. R., Shuler, M. L. & Shain, W. Effects of glial cells on electrode impedance recorded from neural prosthetic devices in vitro. Annals of biomedical engineering 38, 1031–1047 (2010).

48 Woeppel, K., Dhawan, V., Shi, D. & Cui, X. T. Nanotopography-enhanced biomimetic coating maintains bioactivity after weeks of dry storage and improves chronic neural recording. Biomaterials 302, 122326 (2023).

49 Kozai, T. D., Jaquins-Gerstl, A. S., Vazquez, A. L., Michael, A. C. & Cui, X. T. Brain tissue responses to neural implants impact signal sensitivity and intervention strategies. ACS chemical neuroscience 6, 48–67 (2015).

50 Perry, V. H. & Teeling, J. in Seminars in immunopathology. 601–612 (Springer).

51 Salatino, J. W., Ludwig, K. A., Kozai, T. D. Y. & Purcell, E. K. Glial responses to implanted electrodes in the brain. Nature BME (2017).

52 Loane, D. J., Kumar, A., Stoica, B. A., Cabatbat, R. & Faden, A. I. Progressive neurodegeneration after experimental brain trauma: association with chronic microglial activation. Journal of Neuropathology & Experimental Neurology 73, 14–29 (2014).

53 Fiáth, R. et al. Slow insertion of silicon probes improves the quality of acute neuronal recordings. Scientific Reports 9, 111 (2019).

54 Chen, K., Wellman, S. M., Yaxiaer, Y., Eles, J. R. & Kozai, T. D. In vivo spatiotemporal patterns of oligodendrocyte and myelin damage at the neural electrode interface. Biomaterials 268, 120526 (2021).

55 Wellman, S. M., Li, L., Yaxiaer, Y., McNamara, I. & Kozai, T. D. Revealing spatial and temporal patterns of cell death, glial proliferation, and blood-brain barrier dysfunction around implanted intracortical neural interfaces. Frontiers in neuroscience 13, 493 (2019).

56 Yang, Q. et al. Zwitterionic polymer coating suppresses microglial encapsulation to neural implants in vitro and in vivo. Advanced biosystems 4, 1900287 (2020).

57 Wellman, S. M. et al. A materials roadmap to functional neural interface design. Advanced functional materials 28, 1701269 (2018).

58 Yu, K., Niu, X., Krook-Magnuson, E. & He, B. Intrinsic functional neuron-type selectivity of transcranial focused ultrasound neuromodulation. Nature communications 12, 2519 (2021).

59 Niu, X., Yu, K. & He, B. Transcranial focused ultrasound induces sustained synaptic plasticity in rat hippocampus. Brain stimulation 15, 352–359 (2022).

60 Kamimura, H. A., Conti, A., Toschi, N. & Konofagou, E. E. Ultrasound neuromodulation: Mechanisms and the potential of multimodal stimulation for neuronal function assessment. Frontiers in physics 8, 150 (2020).

61 Jiang, X. et al. A review of low-intensity pulsed ultrasound for therapeutic applications. IEEE Transactions on Biomedical Engineering 66, 2704–2718 (2018).

62 Hsu, C. H., Pan, Y. J., Zheng, Y. T., Lo, R. Y. & Yang, F. Y. Ultrasound reduces inflammation by modulating M1/M2 polarization of microglia through STAT1/STAT6/PPARγ signaling pathways. CNS Neuroscience & Therapeutics (2023).

63 Lai, S.-W. et al. Regulatory effects of neuroinflammatory responses through brain-derived neurotrophic factor signaling in microglial cells. Molecular Neurobiology 55, 7487–7499 (2018).

64 Su, W.-S., Wu, C.-H., Song, W.-S., Chen, S.-F. & Yang, F.-Y. Low-intensity pulsed ultrasound ameliorates glia-mediated inflammation and neuronal damage in experimental intracerebral hemorrhage conditions. Journal of Translational Medicine 21, 565 (2023).

65 Rezayat, E. & Toostani, I. G. A review on brain stimulation using low intensity focused ultrasound. Basic and clinical neuroscience 7, 187 (2016).

66 Bystritsky, A. et al. A review of low-intensity focused ultrasound pulsation. Brain stimulation 4, 125–136 (2011).

67 Yue, Z., Moulton, S. E., Cook, M., O’Leary, S. & Wallace, G. G. Controlled delivery for neuro-bionic devices. Advanced Drug Delivery Reviews 65, 559–569 (2013).

68 Kozai, T. D., Eles, J. R., Vazquez, A. L. & Cui, X. T. Two-photon imaging of chronically implanted neural electrodes: Sealing methods and new insights. Journal of neuroscience methods 258, 46–55 (2016).

69 Rueden, C. T. et al. ImageJ2: ImageJ for the next generation of scientific image data. BMC bioinformatics 18, 1–26 (2017).

70 Thevenaz, P., Ruttimann, U. E. & Unser, M. A pyramid approach to subpixel registration based on intensity. IEEE transactions on image processing 7, 27–41 (1998).

71 Kettenmann, H., Hanisch, U.-K., Noda, M. & Verkhratsky, A. Physiology of microglia. Physiological reviews 91, 461–553 (2011).

72 Nahm, F. S. Receiver operating characteristic curve: overview and practical use for clinicians. Korean journal of anesthesiology 75, 25–36 (2022).

73 Onodera, J., Nagata, H., Nakashima, A., Ikegaya, Y. & Koyama, R. Neuronal brain-derived neurotrophic factor manipulates microglial dynamics. Glia 69, 890–904 (2021).

74 Sezgin, M. & Sankur, B. l. Survey over image thresholding techniques and quantitative performance evaluation. Journal of Electronic imaging 13, 146–168 (2004).

75 Li, C. & Tam, P. K.-S. An iterative algorithm for minimum cross entropy thresholding. Pattern recognition letters 19, 771–776 (1998).

76 Li, C. H. & Lee, C. Minimum cross entropy thresholding. Pattern recognition 26, 617–625 (1993).

77 McDowell, K. P., Berthiaume, A. A., Tieu, T., Hartmann, D. A. & Shih, A. Y. VasoMetrics: unbiased spatiotemporal analysis of microvascular diameter in multi-photon imaging applications. Quant Imaging Med Surg 11, 969–982, doi:10.21037/qims-20-920 (2021).

78 Bullitt, E., Aylward, S. R., Van Dyke, T. & Lin, W. Computer-assisted measurement of vessel shape from 3T magnetic resonance angiography of mouse brain. Methods 43, 29–34, doi:10.1016/j.ymeth.2007.03.009 (2007).

79 Wellman, S. M., Cambi, F. & Kozai, T. D. The role of oligodendrocytes and their progenitors on neural interface technology: a novel perspective on tissue regeneration and repair. Biomaterials 183, 200–217 (2018).

80 Ridler, T. & Calvard, S. Picture thresholding using an iterative selection method. IEEE Trans. Syst. Man Cybern 8, 630–632 (1978).

81 Kozai, T. D. Y. et al. Reduction of neurovascular damage resulting from microelectrode insertion into the cerebral cortex using in vivo two-photon mapping. Journal of neural engineering 7, 046011 (2010).

82 Bjornsson, C. et al. Effects of insertion conditions on tissue strain and vascular damage during neuroprosthetic device insertion. Journal of neural engineering 3, 196 (2006).

83 Michelson, N. J. et al. Multi-scale, multi-modal analysis uncovers complex relationship at the brain tissue-implant neural interface: new emphasis on the biological interface. Journal of neural engineering 15, 033001 (2018).

84 Eles, J. R., Vazquez, A. L., Kozai, T. D. & Cui, X. T. In vivo imaging of neuronal calcium during electrode implantation: spatial and temporal mapping of damage and recovery. Biomaterials 174, 79–94 (2018).

85 Madry, C. et al. Microglial ramification, surveillance, and interleukin-1β release are regulated by the two-pore domain K+ channel THIK-1. Neuron 97, 299–312. e296 (2018).

86 Yang, Q., Vazquez, A. L. & Cui, X. T. Long-term in vivo two-photon imaging of the neuroinflammatory response to intracortical implants and micro-vessel disruptions in awake mice. Biomaterials 276, 121060 (2021).

87 Bisht, K. et al. Capillary-associated microglia regulate vascular structure and function through PANX1-P2RY12 coupling in mice. Nature Communications 12, 5289 (2021).

88 Chen, T.-T., Lan, T.-H. & Yang, F.-Y. Low-intensity pulsed ultrasound attenuates LPS-induced neuroinflammation and memory impairment by modulation of TLR4/NF-κB signaling and CREB/BDNF expression. Cerebral Cortex 29, 1430–1438 (2019).

89 Chang, J.-W., Wu, M.-T., Song, W.-S. & Yang, F.-Y. Ultrasound stimulation suppresses LPS-induced proinflammatory responses by regulating NF-κB and CREB activation in microglial cells. Cerebral Cortex 30, 4597–4606 (2020).

90 Davalos, D. et al. Fibrinogen-induced perivascular microglial clustering is required for the development of axonal damage in neuroinflammation. Nature communications 3, 1227 (2012).

91 Davalos, D. et al. ATP mediates rapid microglial response to local brain injury in vivo. Nature neuroscience 8, 752–758 (2005).

92 Lou, N. et al. Purinergic receptor P2RY12-dependent microglial closure of the injured blood–brain barrier. Proceedings of the National Academy of Sciences 113, 1074–1079 (2016).

93 Qin, L., Liu, Y., Hong, J. S. & Crews, F. T. NADPH oxidase and aging drive microglial activation, oxidative stress, and dopaminergic neurodegeneration following systemic LPS administration. Glia 61, 855–868 (2013).

94 Neher, J. J. & Cunningham, C. Priming microglia for innate immune memory in the brain. Trends in Immunology 40, 358–374 (2019).

95 Polikov, V. S., Tresco, P. A. & Reichert, W. M. Response of brain tissue to chronically implanted neural electrodes. Journal of neuroscience methods 148, 1–18 (2005).

96 Williams, J. C., Hippensteel, J. A., Dilgen, J., Shain, W. & Kipke, D. R. Complex impedance spectroscopy for monitoring tissue responses to inserted neural implants. Journal of neural engineering 4, 410 (2007).

97 da Fonseca, A. C. C. et al. The impact of microglial activation on blood-brain barrier in brain diseases. Frontiers in cellular neuroscience 8, 362 (2014).

98 Romano, A. et al. Linking lipid peroxidation and neuropsychiatric disorders: focus on 4-hydroxy-2-nonenal. Free Radical Biology and Medicine 111, 281–293 (2017).

99 Mosher, K. I. & Wyss-Coray, T. Microglial dysfunction in brain aging and Alzheimer’s disease. Biochemical pharmacology 88, 594–604 (2014).

100 Chen, K., Padilla, C. G., Kiselyov, K. & Kozai, T. D. Cell-specific alterations in autophagy-lysosomal activity near the chronically implanted microelectrodes. Biomaterials 302, 122316 (2023).

101 Wang, G. et al. Neuronal accumulation of peroxidated lipids promotes demyelination and neurodegeneration through the activation of the microglial NLRP3 inflammasome. Nature Aging 1, 1024–1037 (2021).

102 Wang, M. et al. Revisiting the intersection of microglial activation and neuroinflammation in Alzheimer’s disease from the perspective of ferroptosis. Chemico-Biological Interactions, 110387 (2023).

103 Filipello, F. et al. Defects in lysosomal function and lipid metabolism in human microglia harboring a TREM2 loss of function mutation. Acta Neuropathologica, 1–24 (2023).

104 Bates, K. A., Martins, R. N. & Harvey, A. R. Oxidative stress in a rat model of chronic gliosis. Neurobiology of Aging 28, 995–1008 (2007).

105 Rayatpour, A., Foolad, F., Heibatollahi, M., Khajeh, K. & Javan, M. Ferroptosis inhibition by deferiprone, attenuates myelin damage and promotes neuroprotection in demyelinated optic nerve. Scientific Reports 12, 19630 (2022).

106 Grandati, M. et al. Calcium-independent NO-synthase activity and nitrites/nitrates production in transient focal cerebral ischaemia in mice. British journal of pharmacology 122, 625 (1997).

107 Kedarasetti, R. T., Drew, P. J. & Costanzo, F. Arterial vasodilation drives convective fluid flow in the brain: a poroelastic model. Fluids and Barriers of the CNS 19, 1–24 (2022).

108 Chen, R. et al. Protective effects of low-intensity pulsed ultrasound (LIPUS) against cerebral ischemic stroke in mice by promoting brain vascular remodeling via the inhibition of ROCK1/p-MLC2 signaling pathway. *Cerebral Cortex*, bhad330 (2023).

109 Hosford, P. S. & Gourine, A. V. What is the key mediator of the neurovascular coupling response? Neuroscience & Biobehavioral Reviews 96, 174–181 (2019).

110 Garcia, V. et al. PIEZO1 expression at the glio-vascular unit adjusts to neuroinflammation in seizure conditions. Neurobiology of Disease, 106297 (2023).

111 De Logu, F. et al. Schwann cell TRPA1 mediates neuroinflammation that sustains macrophage-dependent neuropathic pain in mice. Nature communications 8, 1–16 (2017).

112 Xia, M. et al. TRPA1 activation-induced myelin degradation plays a key role in motor dysfunction after intracerebral hemorrhage. Frontiers in Molecular Neuroscience 12, 98 (2019).

113 Sun, J.-x., et al. Activation of TRPV1 receptor facilitates myelin repair following demyelination via the regulation of microglial function. Acta Pharmacologica Sinica 44, 766–779 (2023).

114 Hu, R., Yang, Z.-y., Li, Y.-h. & Zhou, Z. LIPUS promotes endothelial differentiation and angiogenesis of periodontal ligament stem cells by activating Piezo1. International Journal of Stem Cells 15, 372–383 (2022).

115 Haynes, S. E. et al. The P2Y12 receptor regulates microglial activation by extracellular nucleotides. Nature neuroscience 9, 1512–1519 (2006).

116 Higashi, Y. Lower urinary tract symptoms/benign prostatic hypertrophy and vascular function: Role of the nitric oxide–phosphodiesterase type 5–cyclic guanosine 3′, 5′-monophosphate pathway. International Journal of Urology 24, 412–424 (2017).

117 Larsson, C. Protein kinase C and the regulation of the actin cytoskeleton. Cellular signalling 18, 276–284 (2006).

118 Fu, P.-C. et al. The Rho-associated kinase inhibitors Y27632 and fasudil promote microglial migration in the spinal cord via the ERK signaling pathway. Neural Regeneration Research 13, 677 (2018).

119 Liu, Y.-J. et al. Late endosomes promote microglia migration via cytosolic translocation of immature protease cathD. Science Advances 6, eaba5783 (2020).

120 Wu, F. et al. CXCR2 antagonist attenuates neutrophil transmigration into brain in a murine model of LPS induced neuroinflammation. Biochemical and Biophysical Research Communications 529, 839–845 (2020).

121 Lu, H. et al. Identification of genes responsive to low-intensity pulsed ultrasound stimulations. Biochemical and biophysical research communications 378, 569–573 (2009).

122 Dou, Y. et al. Microglial migration mediated by ATP-induced ATP release from lysosomes. Cell research 22, 1022–1033 (2012).

123 Mastorakos, P. et al. Temporally distinct myeloid cell responses mediate damage and repair after cerebrovascular injury. Nature neuroscience 24, 245–258 (2021).

124 Lam, D. V. et al. Platelets and hemostatic proteins are co-localized with chronic neuroinflammation surrounding implanted intracortical microelectrodes. Acta Biomaterialia (2023).

125 Adams, R. A. et al. The fibrin-derived γ377-395 peptide inhibits microglia activation and suppresses relapsing paralysis in central nervous system autoimmune disease. The Journal of experimental medicine 204, 571–582 (2007).

126 Ryu, J. K. et al. Fibrin-targeting immunotherapy protects against neuroinflammation and neurodegeneration. Nature immunology 19, 1212–1223 (2018).

127 Baik, S. H. et al. A breakdown in metabolic reprogramming causes microglia dysfunction in Alzheimer’s disease. Cell metabolism 30, 493–507. e496 (2019).

128 Hu, Y. et al. Replicative senescence dictates the emergence of disease-associated microglia and contributes to Aβ pathology. Cell reports 35 (2021).

129 Kaloss, A. M. et al. Noninvasive Low-Intensity Focused Ultrasound Mediates Tissue Protection following Ischemic Stroke. BME Frontiers 2022 (2022).

130 Leng, S. et al. Ion channel Piezo1 activation promotes aerobic glycolysis in macrophages. Frontiers in Immunology 13, 976482 (2022).

131 Chen, J. et al. Potential Application of Low-Intensity Pulsed Ultrasound in Delaying Aging for Mice. Gerontology 68, 558–570 (2022).

132 Chen, F. R., Manzi, J. E., Mehta, N., Gulati, A. & Jones, M. A Review of Laser Therapy and Low-Intensity Ultrasound for Chronic Pain States. Current Pain and Headache Reports 26, 57–63 (2022).

133 Ichijo, S. et al. Low-intensity pulsed ultrasound therapy promotes recovery from stroke by enhancing angio-neurogenesis in mice in vivo. Scientific reports 11, 4958 (2021).

134 dos Santos Tramontin, N., et al. Effects of Low-Intensity Transcranial Pulsed Ultrasound Treatment in a Model of Alzheimer’s Disease. Ultrasound in Medicine & Biology 47, 2646–2656 (2021).

135 Zhou, H. et al. Wearable ultrasound improves motor function in an MPTP mouse model of Parkinson’s disease. IEEE Transactions on Biomedical Engineering 66, 3006–3013 (2019).

136 Yang, F.-Y., Huang, L.-H., Wu, M.-T. & Pan, Z.-Y. Ultrasound Neuromodulation Reduces Demyelination in a Rat Model of Multiple Sclerosis. International Journal of Molecular Sciences 23, 10034 (2022).

137 Jia, J. et al. The role of microglial phagocytosis in ischemic stroke. Frontiers in Immunology 12, 790201 (2022).

138 Eyo, U. & Dailey, M. E. Effects of oxygen-glucose deprivation on microglial mobility and viability in developing mouse hippocampal tissues. Glia 60, 1747–1760 (2012).

139 Neumann, J. et al. Microglia provide neuroprotection after ischemia. The FASEB journal 20, 714–716 (2006).

140 Masuda, T., Croom, D., Hida, H. & Kirov, S. A. Capillary blood flow around microglial somata determines dynamics of microglial processes in ischemic conditions. Glia 59, 1744–1753 (2011).

141 van Rossum, D. & Hanisch, U.-K. Microglia. Metabolic brain disease 19, 393–411 (2004).

142 Nimmerjahn, A., Kirchhoff, F. & Helmchen, F. Resting microglial cells are highly dynamic surveillants of brain parenchyma in vivo. Science 308, 1314–1318 (2005).

143 Fetler, L. & Amigorena, S. Brain under surveillance: the microglia patrol. Science 309, 392–393 (2005).

144 Tao, L. et al. Microglia modulation with 1070-nm light attenuates Aβ burden and cognitive impairment in Alzheimer’s disease mouse model. Light: Science & Applications 10, 179 (2021).

145 Gao, H.-M. et al. Neuroinflammation and α-synuclein dysfunction potentiate each other, driving chronic progression of neurodegeneration in a mouse model of Parkinson’s disease. Environmental health perspectives 119, 807–814 (2011).

146 Choi, J. H. et al. Age-related changes in ionized calcium-binding adapter molecule 1 immunoreactivity and protein level in the gerbil hippocampal CA1 region. Journal of Veterinary Medical Science 69, 1131–1136 (2007).

147 Kubanek, J., Shukla, P., Das, A., Baccus, S. A. & Goodman, M. B. Ultrasound elicits behavioral responses through mechanical effects on neurons and ion channels in a simple nervous system. Journal of Neuroscience 38, 3081–3091 (2018).

148 Oh, S.-J. et al. Ultrasonic neuromodulation via astrocytic TRPA1. Current Biology 29, 3386–3401. e3388 (2019).

149 Blackmore, D. G., Razansky, D. & Götz, J. Ultrasound as a versatile tool for short-and long-term improvement and monitoring of brain function. Neuron 111, 1174–1190 (2023).

150 Darrow, D. P., O’Brien, P., Richner, T. J., Netoff, T. I. & Ebbini, E. S. Reversible neuroinhibition by focused ultrasound is mediated by a thermal mechanism. Brain stimulation 12, 1439–1447 (2019).

151 Yuan, Y., Wang, Z., Liu, M. & Shoham, S. Cortical hemodynamic responses induced by low-intensity transcranial ultrasound stimulation of mouse cortex. NeuroImage 211, 116597 (2020).

152 Mesiwala, A. H. et al. High-intensity focused ultrasound selectively disrupts the blood-brain barrier in vivo. Ultrasound in medicine & biology 28, 389–400 (2002).

153 Robertson, J. L., Cox, B. T., Jaros, J. & Treeby, B. E. Accurate simulation of transcranial ultrasound propagation for ultrasonic neuromodulation and stimulation. The Journal of the Acoustical Society of America 141, 1726–1738 (2017).

154 Guo, J., Song, X., Chen, X., Xu, M. & Ming, D. Mathematical model of ultrasound attenuation with skull thickness for transcranial-focused ultrasound. Frontiers in Neuroscience 15, 778616 (2022).

155 Adams, M. S., Lotz, J. C. & Diederich, C. J. In silico feasibility assessment of extracorporeal delivery of low-intensity pulsed ultrasound to intervertebral discs within the lumbar spine. Physics in Medicine & Biology 65, 215011 (2020).

